# Small effect size leads to reproducibility failure in resting-state fMRI studies

**DOI:** 10.1101/285171

**Authors:** Xi-Ze Jia, Na Zhao, Barek Barton, Roxana Burciu, Nicolas Carrière, Antonio Cerasa, Bo-Yu Chen, Jun Chen, Stephen Coombes, Luc Defebvre, Christine Delmaire, Kathy Dujardin, Fabrizio Esposito, Guo-Guang Fan, Di Nardo Federica, Yi-Xuan Feng, Brett W. Fling, Saurabh Garg, Moran Gilat, Martin Gorges, Shu-Leong Ho, Fay B. Horak, Xiao Hu, Xiao-Fei Hu, Biao Huang, Pei-Yu Huang, Ze-Juan Jia, Christy Jones, Jan Kassubek, Lenka Krajcovicova, Ajay Kurani, Jing Li, Qian Li, Ai-Ping Liu, Bo Liu, Hu Liu, Wei-Guo Liu, Renaud Lopes, Yu-Ting Lou, Wei Luo, Tara Madhyastha, Ni-Ni Mao, Grainne McAlonan, Martin J. McKeown, Shirley YY Pang, Aldo Quattrone, Irena Rektorova, Alessia Sarica, Hui-Fang Shang, James Shine, Priyank Shukla, Tomas Slavicek, Xiao-Peng Song, Gioacchino Tedeschi, Alessandro Tessitore, David Vaillancourt, Jian Wang, Jue Wang, Z. Jane Wang, Lu-Qing Wei, Xia Wu, Xiao-Jun Xu, Lei Yan, Jing Yang, Wan-Qun Yang, Nai-Lin Yao, De-Long Zhang, Jiu-Quan Zhang, Min-Ming Zhang, Yan-Ling Zhang, Cai-Hong Zhou, Chao-Gan Yan, Xi-Nian Zuo, Mark Hallett, Tao Wu, Yu-Feng Zang

**Author notes:** Correspondence should be addressed to Y.F.Z. (;).

## Abstract

Thousands of papers using resting-state functional magnetic resonance imaging (RS-fMRI) have been published on brain disorders. Results in each paper may have survived correction for multiple comparison. However, since there have been no robust results from large scale meta-analysis, we do not know how many of published results are truly positives. The present meta-analytic work included 60 original studies, with 57 studies (4 datasets, 2266 participants) that used a between-group design and 3 studies (1 dataset, 107 participants) that employed a within-group design. To evaluate the effect size of brain disorders, a very large neuroimaging dataset ranging from neurological to psychiatric disorders together with healthy individuals have been analyzed. Parkinson’s disease off levodopa (PD-off) included 687 participants from 15 studies. PD on levodopa (PD-on) included 261 participants from 9 studies. Autism spectrum disorder (ASD) included 958 participants from 27 studies. The meta-analyses of a metric named amplitude of low frequency fluctuation (ALFF) showed that the effect size (Hedges’ *g*) was 0.19 - 0.39 for the 4 datasets using between-group design and 0.46 for the dataset using within-group design. The effect size of PD-off, PD-on and ASD were 0.23, 0.39, and 0.19, respectively. Using the meta-analysis results as the robust results, the between-group design results of each study showed high false negative rates (median 99%), high false discovery rates (median 86%), and low accuracy (median 1%), regardless of whether stringent or liberal multiple comparison correction was used. The findings were similar for 4 RS-fMRI metrics including ALFF, regional homogeneity, and degree centrality, as well as for another widely used RS-fMRI metric namely seed-based functional connectivity. These observations suggest that multiple comparison correction does not control for false discoveries across multiple studies when the effect sizes are relatively small. Meta-analysis on un-thresholded *t*-maps is critical for the recovery of ground truth. We recommend that to achieve high reproducibility through meta-analysis, the neuroimaging research field should share raw data or, at minimum, provide un-thresholded statistical images.

## Introduction

Meta-analysis is the process of combining the results of independently conducted studies^1^. Resting-state functional magnetic resonance imaging (RS-fMRI), a technique to examine the spontaneous brain activity^2^ non-invasively with high spatial and temporal resolution, is available in most hospitals, and easy for patients to tolerate. Thousands of papers using RS-fMRI for research on brain disorders have been published since the seminal RS-fMRI study of Biswal and colleagues 23 years ago^3^. Meta-analysis can provide robust results of brain disorders. However, few high-quality meta-analytical papers using RS-fMRI to study brain disorders have been published on account of two main reasons.

First, there are countless analytical methods and options. For example, seed-based functional connectivity is the most widely used method in RS-fMRI studies. However, the locations of seed-regions vary substantially from study to study^4,5^; therefore, it is impossible to perform high-quality meta-analysis for a fixed seed location. While various analytical methods may help investigate the complexity of human brain function from different angles, few papers have used very similar analytical methods. Hence, it is difficult for a meta-analysis to include an adequate number of original studies. To date, we have found only 7 RS-fMRI meta-analysis papers^6-12^ in which the original studies used highly comparable analytical methods.

Second, almost all published papers have only reported a limited number of brain regions that survived multiple comparison correction (MCC). This means that the effect size for all other voxels was essentially reported as zero, and hence, it is highly likely that a “positive or zero” bias will occur in subsequent meta-analysis. The effect size remains unknown for most RS-fMRI studies of brain disorders. If the effect size is not large enough, some reported brain areas are very likely to be false positives and, meanwhile, some true positive brain regions may not survive the multiple comparison correction.

Here, we performed a high-quality RS-fMRI meta-analysis to approximate the effect size of 60 studies from 5 datasets (total N = 2373). To evaluate the effect size of brain disorders, a very large neuroimaging dataset ranging from neurological (PD) to psychiatric disorders (ASD) together with healthy individuals were included. To avoid the issue of “positive or zero” bias, we performed the meta-analysis on unthresholded *t*-maps. We evaluated four metrics, including three whole-brain voxel-wise metrics, i.e., amplitude of low frequency fluctuation (ALFF)^13^, regional homogeneity (ReHo)^14^, degree centrality (DC)^15,16^ and a mostly used functional connectivity metric, i.e., seed-based functional connectivity (SFC)^3^.

## Results

The 5 datasets were Parkinson’s disease off levodopa (PD-off) vs. healthy controls (HC), PD on levodopa (PD-on) vs. HC, autism spectrum disorder (ASD) vs HC, healthy male vs. female (MF), and eyes open vs. eyes closed (EOEC) (**Table 1**). The details are listed as the supplementary information (**Supplementary Note 1**).

**Table 1.** Information for each dataset. PD-off, Parkinson’s disease off levodopa vs. healthy controls (HC). PD-on, PD on levodopa vs. HC. ASD, autism spectrum disorder vs. HC. MF, healthy male vs. female. EOEC, eye open vs. eyes closed.

|  | Dataset | Number<br>of studies | N |
| --- | --- | --- | --- |
| Between-group design | PD-off | 15 | 687 |
|  | PD-on | 9 | 261 |
|  | ASD | 27 | 958 |
|  | MF | 6 | 360 |
| Within-group design | EOEC | 3 | 107 |
|  |  | 60 | 2373 |

### Effect size of each dataset

For the 4 datasets of between-group design, the effect size of the meta-analysis was very small overall (**Fig. 1**), and only a small number of voxels had an effect size of > 0.5. For example, for a typical ALFF metric, the median effect size of the PD-off, PD-on, ASD, and MF was 0.19 - 0.39. For the within-group design, EOEC showed a higher median effect size (0.46) (**Fig. 1a**). The most widely used RS-fMRI metric, i.e., seed-based functional connectivity (SFC), showed similar results (**Fig. 1b**), as did the other 2 metrics, i.e., ReHo and DC (**Supplementary Fig. 5**).

**Figure 1.**
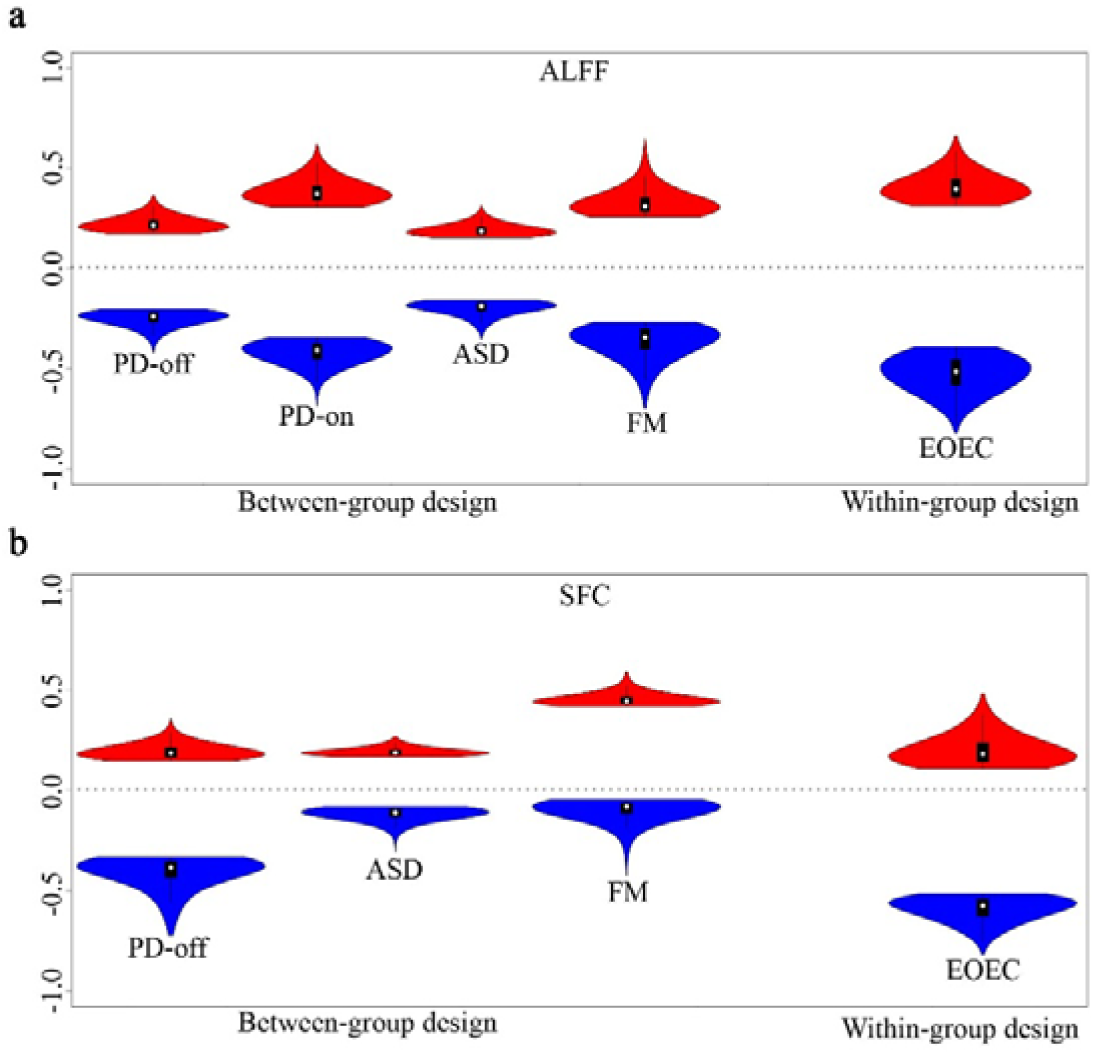
Effect size (Hedges’ g) of meta-analyses for ALFF (a) and SFC (b). Red indicates higher ALFF or SFC in PD (PD-off and PD-on) than controls, ASD than control, males than females, and EO than EC, respectively. Blue indicates the opposite. The white dot in the violin plot indicates the median, and the black bar indicates the interquartile range. SFC was not analyzed for PD-off and PD-on datasets (Please see **Supplementary Note 1**). The plot was created by R package *vioplot* 0.2 (https://cran.r-project.org/web/packages/vioplot/index.html). PD-off, Parkinson’s disease off levodopa vs. healthy controls (HC). PD-on, PD on levodopa vs. HC. ASD, autism spectrum disorder vs HC. MF, healthy male vs. female. EOEC, eye open vs. eyes closed. ALFF, amplitude of low frequency fluctuation. SFC, seed-based functional connectivity.

### High false negative rate and high false discovery rate

Using meta-analytical results as the robust result, we calculated the false negative rate (FNR), accuracy, and false discovery rate (FDR) for each study in each dataset. We used 3 methods for multiple comparison correction, including permutation test with threshold-free cluster enhancement (TFCE) (corrected *P* < 0.05)^17^, false discovery rate correction (FDR-c, *q* < 0.05)^18^, and an arbitrary threshold (Arbi, individual voxel *P* < 0.01 and cluster size > 50 voxels). As shown in **Fig. 2a-c** of ALFF results, all between-group design studies showed very high FNR and very low accuracy. The within-group design, i.e., EOEC, showed better performance than the between-group design. The above results were similar for SFC (**Fig. 2d-f**), ReHo, and DC (**Supplementary Fig. 6 - 10**).

**Figure 2.**
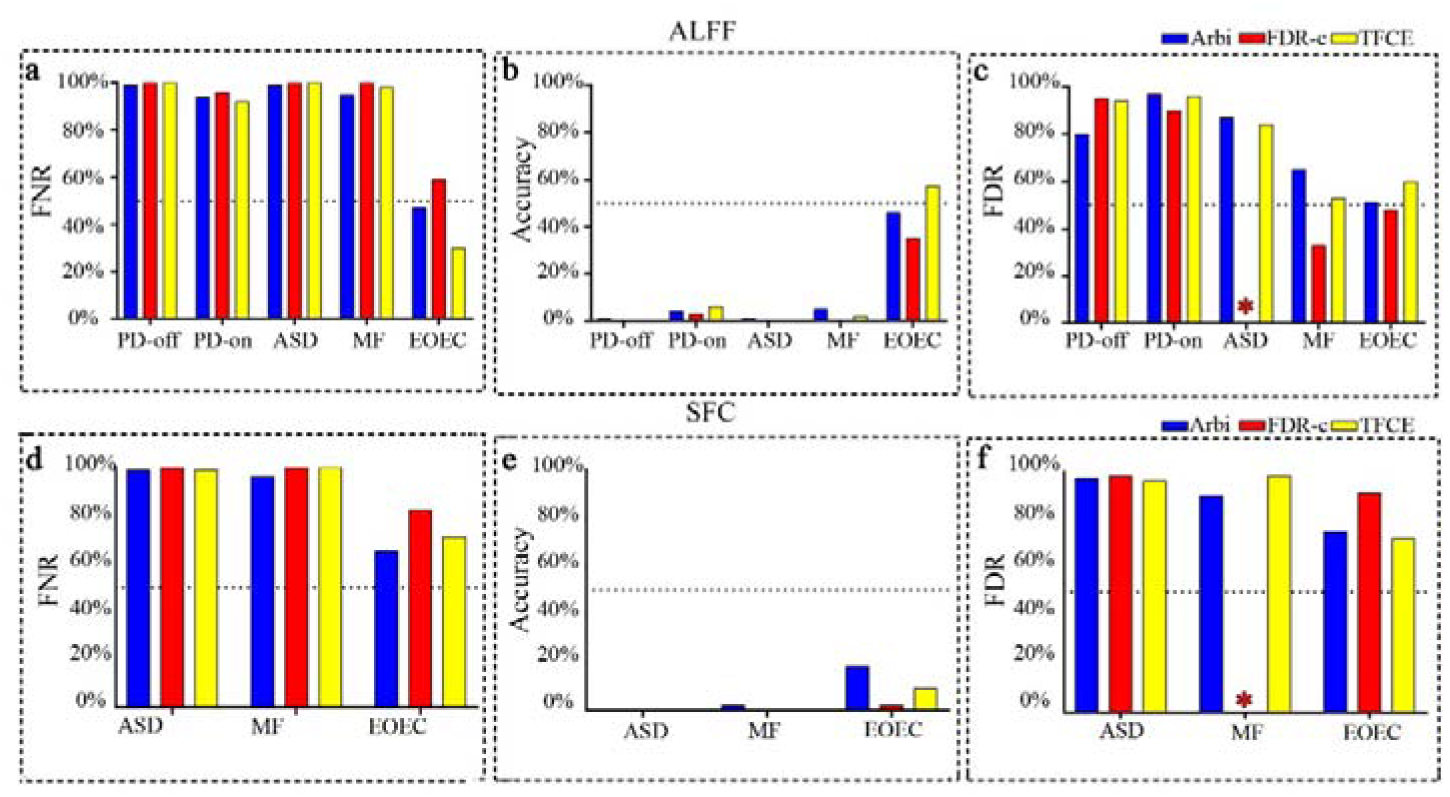
FNR, accuracy, and FDR of ALFF and SFC. Using meta-analytical results as the robust result, the mean FNR **(a, d)**, accuracy **(b, e)**, and FDR **(c, f)** were calculated from thresholded *t*-images of all studies for each dataset. SFC was not analyzed for PD-off and PD-on datasets (Please see **Supplementary Note 1**). GraphPad Prism 7.00 was used for plotting (http://www.graphpad.com). *: No voxels survived FDR-c for any study of ASD **(c)** and MF **(f)**. PD-off, Parkinson’s disease off levodopa vs. healthy controls (HC). FNR, false negative rate. FDR, false discovery rate. PD-on, PD on levodopa vs. HC. ASD, autism spectrum disorder vs HC. MF, healthy male vs. female. EOEC, eye open vs. eyes closed. ALFF, amplitude of low frequency fluctuation. SFC, seed-based functional connectivity. Arbi, arbitrary threshold. FDR-c, false discovery rate correction. TFCE, threshold-free cluster enhancement.

### Exemplar brain regions of meta-analysis

We took ALFF as an exemplar metric to show the pattern of differences for a few datasets. Significant differences were found between PD patients and controls in the bilateral putamen, bilateral sensorimotor cortices, and other areas (**Fig. 3**). Patients with ASD showed decreased ALFF in the posterior cingulate cortex (PCC) and increased ALFF in the cerebellum and frontal cortex. EC showed higher ALFF than EO in the sensorimotor cortex and lower ALFF in the lateral occipital cortex. All detailed differences in brain patterns were shown in **Supplementary Fig. 3** (between-group design) and **Supplementary Fig. 4** (within-group design). All resultant un-thresholded statistical images can be downloaded at https://github.com/jiaxize/MetaAnalysisPaper.

**Figure 3.**
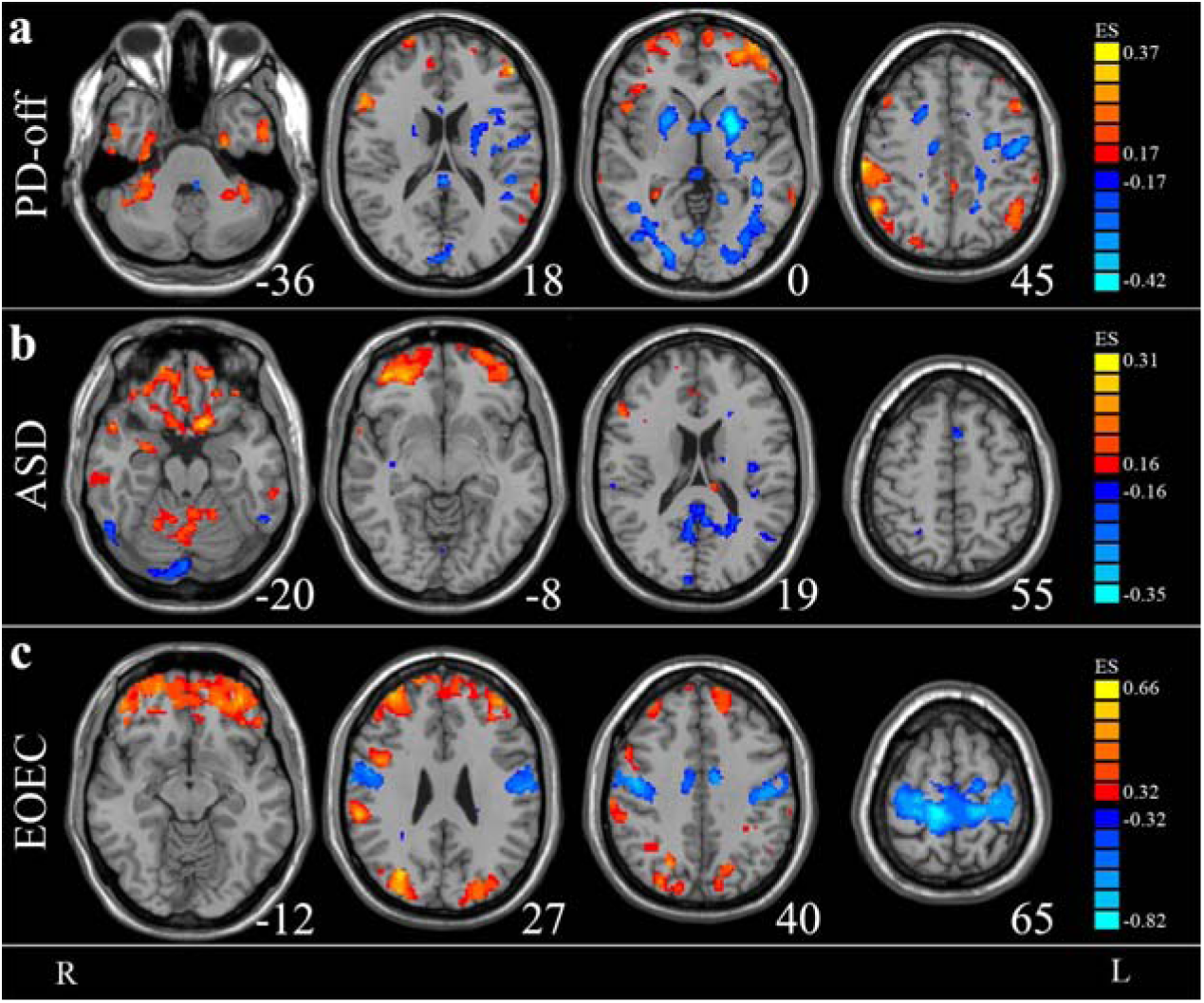
Exemplar brain regions of meta-analyses of ALFF. Warm colors indicate higher ALFF in PD than control, ASD than control, and EO than EC. Cold colors indicate the opposite. **(a)** The most evident finding in the PD group during OFF-dopaminergic phase was the presence of lower ALFF in the bilateral putamen and pallidum; **(b)** High ALFF in the orbitofrontal cortex critically involved in pathophysiological mechanisms of ASD; **(c)** The results of EOEC data revealed higher ALFF in the sensorimotor cortex and lower ALFF in the lateral occipital cortex in EC than in EO. A combination threshold *P* < 0.005, *z* > 1, and cluster size > 10 voxels was used. L: left side of the brain. R: right side of the brain. ES: Effect size (Hedges’ *g*). PD-off, Parkinson’s disease off levodopa vs. healthy controls (HC). ASD, autism spectrum disorder vs HC. EOEC, eye open vs. eyes closed.

## Discussion

### Possible reasons for high FNR and FDR

Regardless of stringent (i.e., FDR-c and TFCE) or liberal (i.e., Arbi) thresholds for multiple comparison correction, we found a very high FNR and FDR for each study, especially for studies of brain disorders (**Fig. 2a-f**). The overall results were similar with or without accounting for head motion parameters or global mean time course. The high FNR could be explained by the small effect size because most voxels will not survive an uncorrected threshold of *P* < 0.05. The high FDR could also be due to the high dimensionality (tens of thousands of voxels). The high FNR and high FDR could be among the most important explanations for low reproducibility in RS-fMRI studies of brain disorders. This concern has frequently been addressed in biomedical research^19-26^.

In the present study, we analyzed 60 studies from multiple research centers and provided strong empirical evidence that the results of a single RS-fMRI study with between-group design have high risk for FNR and FDR.

### Despite the small effect size for RS-fMRI, the results of image-based meta-analyses revealed some convergent results

PD-off meta-analysis showed decreased ALFF in the bilateral putamen (**Fig. 3**), a key region which has been well-documented in previous dopaminergic positron emission tomography (PET)^27^ and meta-analysis of RS-fMRI^11^. The results of EOEC data were very consistent with those of previous RS-fMRI studies^28,29^, e.g., higher ALFF in the sensorimotor cortex and lower ALFF in the lateral occipital cortex in EC than in EO (**Fig. 3**). The consistent results of EOEC are probably attributable to relatively larger effect size of EOEC data (**Fig. 1**). The ABIDE initiative provided a large-scale dataset comprised of data from over 24 international brain imaging laboratories (http://fcon_1000.projects.nitrc.org/indi/abide). However, few studies have reported large-scale meta-analytical results based on whole-brain voxel-wise (WBVW) analytical metrics of RS-fMRI. ALFF, ReHo, and DC are three widely used WBVW analytical metrics of RS-fMRI data. These metrics are more suitable for meta-analysis^4^. The current meta-analysis found convergent abnormal brain activity in autism patients in a few brain regions, e.g., the PCC (**Fig. 3**). These brain regions could be used to inform *a priori* hypothesis of the location of an expected effect in future autism neuroimaging studies.

In the RS-fMRI community, some multi-center and large-scale raw data datasets of health adults are publicly available (e.g., http://fcon_1000.projects.nitrc.org, https://www.humanconnectome.org/). However, very limited number of brain disorder datasets have been made publicly available. ABIDE provided a large open access dataset of ASD, but, few studies have performed large-scale image-based meta-analysis using WBVW metrics. Despite the small effect size and variations in clinical features from site to site for PD and ASD patients, meta-analysis can provide valuable robust results from multi-center datasets and would help precisely localize the abnormal brain activity common to a specific disorder beyond subtype for comorbidity.

### Limitations and future directions

First, we examined only three WBVW RS-fMRI metrics and one functional connectivity metric, i.e., seed-based functional connectivity analysis. Thus, future studies should investigate whether the magnitude of the effect size is similarly small for other metrics. Further, based on large, multicenter datasets or strong hypotheses regarding a specific brain region or network, meta-analysis could help researchers to discover more robust results. Based robust results (e.g., putamen for PD), methodology studies could provide new analytical methods which could yield larger effect sizes. Second, for the patient datasets of PD and ASD, we did not take the disease subtype or symptoms severity into account. Future studies should investigate the subtype- or symptom-specific abnormalities. Third, sharing raw data is critical for large-scale image-based meta-analysis. It has been proposed that the neuroimaging research field should develop an effective mechanism for sharing image data^30^. Given the current social and technical difficulties in sharing data, publicly sharing the unthresholded statistical maps may be more feasible in most instances. WBVW analytical methods are more suitable for meta-analysis^4^. Therefore, we suggest that future RS-fMRI studies should publish their unthresholded statistical images with WBVW analysis, even though the WBVW analysis is not the main focus of a given study.

## ACKNOWLEDGMENTS

This work was supported by the Natural Science Foundation of China (No. 81520108016, 81661148045, and 31471084 to YFZ) and other grants (Please see **Supplementary Note 2**). Dr. Zang is partially supported by the “Qian Jiang Distinguished Professor” program.

## AUTHOR CONTRIBUTIONS

X.Z.J., D.V., C.G.Y., X.N.Z., M.H., T.W. and Y.F.Z. designed the research; X.Z.J., N.Z., B.B., M.B., R.B., N.C., A.C., B.Y.C., J.C., L.D., C.D., K.D., F.E., G.G.F., Y.X.F., B.W.F., S.G., Moran G., Martin G., S.L.H., F.B.H., X.H., X.F.H., B.H., P.Y.H., Z.J.J., C.J., J.K., L.K., A.K., J.L., Q.L., A.P.L., B.L., H.L., W.G.L., R.L., Y.T.L., W.L., T.M., N.N.M., G.M., M.J.M., S.Y.Y.P., A.Q., I.R., A.S., H.F.S., J.S., T.S., X.P.S., G.T., A.T., D.V., Jian W., Jue W., Z. J.W., L.Q.W., X.W., X.J.X., L.Y., J.Y., W.Q.Y., N.L.Y., D.L.Z., J.Q.Z., M.M.Z., Y.L.Z, C.H.Z., C.G.Y., X.N.Z., M.H., T.W. and Y.F.Z. collected the data; X.Z.J., N.Z., A.C., D.N.F., Y.X.F., M.G., Z.J.J., J.S., P.S., Jue W. and J.Y. analyzed the data. X.Z.J. and Y.F.Z wrote the first draft of the manuscript, which R.B., S.C., A.C., F.E. A.K., J.K., M.G., G.M.M., T.M.M., J.M.S., A.T., D.V., C.G.Y., and X.N.Z., revised. All authors approved.

## ONLINE METHODS

### Datasets

There were 5 datasets in the current study: PD-off, PD-on, ASD, MF, and EOEC (See **Supplementary Note 1** for details). Some participants were excluded (**Supplementary Table 1 - 4**). The remaining data were summarized in **Supplementary Table 5 - 9**.

### Preprocessing

All preprocessing was performed by RESTplus V1.2 (http://www.restfmri.net/forum/RESTplusV1.2). RESTplus was based on SPM12 (http://www.fil.ion.ucl.ac.uk/spm/software/spm12) and REST V1.8 (http://www.restfmri.net/forum/REST_V1.8). Preprocessing steps included:

1. Discarding the first 10 timepoints.
2. Slice timing correction.
3. Head motion correction.
4. Spatial normalization. Briefly, individual T1-weighted image was co-registered to the mean functional image and then segmented into gray matter (GM), white matter (WM), and cerebrospinal fluid (CSF). Using y_*.nii, all EPI images were spatially normalized to the Montreal Neurological Institute (MNI) space and voxel size was resampled to 3 × 3 × 3 mm^3^, for all datasets except PD datasets. Because of ethical issues, the raw data of most studies of PD-off dataset and PD-on dataset was not sent to Hangzhou Normal University. Instead, the data was analyzed in each research center by using the pipeline (rest_metabatch.m). The resultant images and relevant information (e.g., head motion parameters, spatial normalization, and log files) were sent to Hangzhou Normal University. To reduce the probability of errors during spatial normalization, the RS-fMRI was spatially normalized to EPI template. The amplitude of low frequency fluctuation (ALFF), regional homogeneity (ReHo), and degree centrality (DC) images were sent to Hangzhou Normal University and were analyzed together.
5. Spatial smoothing with an isotropic Gaussian kernel with an FWHM of 6 mm before ALFF and seed-based functional connectivity (SFC) calculation. But for the ReHo and DC images, smoothing was performed after ReHo or DC calculation because smooth can artificially increase the localized similarity of the original time.
6. Removing the linear trend within the time course.
7. Regressing out covariates. The results in the main content of the article were without regressing out of covariates. The supporting information included results of both with and without regressing out covariates. We had two sets of covariates. One set consisted of Friston-24 head motion parameters^31^, white matter (WM) signal, and cerebrospinal fluid (CSF) signal. Another set of parameters included global mean time course, WM signal, and CSF signal.
8. Band-pass filtering (0.01-0.08Hz, for ReHo, DC and SFC).

### Calculation of metrics

ALFF calculation was based on fast Fourier transform (FFT)^13^. Using FFT, each time course was converted to frequency domain. Then, the square root of power spectrum at each frequency was averaged across a specific frequency band (0.01-0.08 in current study). This averaged square root was taken as ALFF. ALFF represents the strength of low frequency oscillations. The ALFF maps were *Z*-standardized, i.e., subtracting the mean value of entire brain and divided by the corresponding standard deviation.

ReHo measures the local synchronization of spontaneous brain activities of neighboring voxels^14^. The Kendall’s coefficient of concordance^32^ of the time course of every 27 nearest neighboring voxels was calculated. A larger ReHo value for a given voxel means the higher regional coherence. ReHo maps were *Z*-standardized as did for ALFF maps.

DC is the total number or weighted sum of significant connections of a voxel with all voxels^15,33^. We calculated the weighted sum of positive correlations using a threshold *r* > 0.2. Z-standardized DC maps were created as did for ALFF and ReHo.

Due to ethical issue, most of the PD datasets were analyzed in each research center (Please see **Supplementary Note 1** for details). And because the SFC was a post-hoc analysis, SFC analyses were performed for ASD, FM, EOEC, but not for PD-off and PD-on. The seed-ROIs were listed **Supplementary Table 10**.

### T-tests and multiple comparison corrections for each study

DPABI V2.3^34^ was used for *t*-tests. Two-sample *t*-tests were performed for studies of between-group design datasets (PD-off, PD-on, ASD, and FM datasets). And paired *t*-tests were performed for studies of within-group design (EOEC dataset).

Three multiple comparison methods were used. One was an arbitrary threshold (Arbi, individual voxel *P* < 0.01, cluster size > 50 voxels and edge connected). The second one was false discovery rate correction (FDR-c, *q* < 0.05)^18^. The Arbi and FDR-c were performed by RESTplus V1.2. The third one, threshold-free cluster enhancement (TFCE)^17^, was performed by DPABI V2.3. The TFCE of DPABI V2.3 was based on integrating PALM package^35^. TFCE used the following parameters: no acceleration, 5000 permutation tests, FWER-correction *P* < 0.05, and two tailed.

### Meta-analysis and FNR, accuracy and FDR

Anisotropic effect size signed differential mapping (AES-SDM 5.141)^36-38^ was used to perform meta-analysis on the unthresholded *t*-maps for all the studies in each dataset. AES-SDM assigns each voxel a measure of effect size, namely the standardized mean difference, known as Hedge’s δ^39^. AES-SDM used the random-effects model to combine the statistical parametric maps. A recommended threshold *P* < 0.005, *z* > 1, cluster size > 10 voxels was used^37^.

We took the meta-analytical results of each dataset as robust result. Then, the false negative rate (*FNR*) was defined as

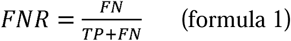

The *FN* is the number of false negative voxels. The *TP* is the number of true positive voxels. The false discovery rate (*FDR*) was defined as

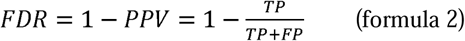

For each dataset, *FDR* was calculated on those studies which have voxels survived the correction. If a study has no any voxel surviving the correction, *FDR* was not computed for that study. The positive predictive value (*PPV*) is the probability that a positive finding reflects a true effect^25^. The *FP* is the number of false positive voxels.

The accuracy is defined as,

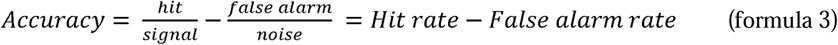

The *hit* is equal to *TP*. The false alarm is equal to *FP. Signal* is the number of voxels of true positive in the robust result. *Noise* is the number of voxels outside the robust result, i.e., true negative.

It should be noted that the voxel size in the results of AES-SDM software was 2 × 2 × 2 mm^3^, but the voxel size of other images was 3 × 3 × 3 mm^3^. So we re-sampled the AES-SDM voxel size into 3 × 3 × 3 mm^3^ by using nearest neighboring method in RESTplus.

