## Supplementary Materials for "Small effect size leads to reproducibility failure in resting-state fMRI studies"

[13] Zhejiang Provincial Key Lab of Ophthalmology.

[14] Department of Health and Exercise Science, Colorado State University.

[15] Pacific Parkinson's Research Centre, University of British Columbia.

[16] Department of Medicine (Neurology) University of British Columbia.

[17] Brain and Mind Center, The University of Sydney.

[18] Department of Neurology, Ulm University.

[19] Division of Neurology, Dept of Medicine, Queen Mary Hospital, University of Hong Kong.
[20] Department of Neurology, School of Medicine, Oregon Health & Science University.
[21] VA Portland Health Care System.
[22] Department of Radiology, The Affiliated Brain Hospital With Nanjing Medical University.
[23] Department of Radiology, Southwest Hospital, Third Military Medical University.
[24] Department of Radiology, Guangdong Academy of Medical Sciences.
[25] Department of Radiology, The Second Affiliated Hospital, Zhejiang University School of Medicine.
[26] Graduate School of Hebei Medical University.
[27] Department of Radiology, Northwestern University Feinberg School of Medicine.
[28] Department of Neurology, Southwest Hospital, Third Military Medical University.
[29] College of Information Science and Technology, Beijing Normal University.
[30] Department of Electrical and Computer Engineering, University of British Columbia.
[31] Department of Neurology, The Affiliated Brain Hospital With Nanjing Medical University.
[32] Department of Pediatrics, The Second Affiliated Hospital, Zhejiang University School of Medicine.
[33] Department of Neurology, The Second Affiliated Hospital, Zhejiang University School of Medicine.
[34] Department of Radiology, University of Washington.
[35] The Sackler Institute for Translational Neuroscience, King's College London and the Biomedical Research Centre for Mental Health at South London and Maudsley NHS Foundation Trust and Institute of Psychiatry, Psychology and Neuroscience, King's College London.
[36] State Key Laboratory for Cognitive Sciences, The University of Hong Kong.
[37] Department of Forensic and Neurodevelopmental Science, Institute of Psychiatry, King's College London.
[38] Institute of Neurology, University "Magna Graecia".
[39] Department of Neurology, West China Hospital, SiChuan University.
[40] MIND Research Network.
[31] Department of Neurology, The Affiliated Brain Hospital With Nanjing Medical University.
[41] Key Laboratory of Exercise and Health Sciences, Ministry of Education, Shanghai University of Sport.
[43] CAS Key Laboratory of Behavioral Science, Institute of Psychology.
[44] University of Chinese Academy of Sciences, Beijing Institute of Geriatrics.
[45] Human Motor Control Section, National Institute of Neurological Disorders and Stroke, National Institutes of Health.
[46] Department of Neurobiology, Neurology and Geriatrics, Xuanwu Hospital of Capital Medical University, Beijing Institute of Geriatrics.
[47] Clinical Center for Parkinson’s Disease, Capital Medical University.
[48] Beijing Key Laboratory for Parkinson’s Disease, Parkinson Disease Center of Beijing Institute for Brain Disorders.
[49] National Clinical Research Center for Geriatric Disorders.

| | |
|---|---|
| Supplementary Note 1 | Information of each dataset |
| Supplementary Note 2 | ACKNOWLEDGMENTS |
| Supplementary Table 1 | Detailed exclusion information for PD-off dataset. |
| Supplementary Table 2 | Detailed exclusion information for PD-on dataset. |
| Supplementary Table 3 | Detailed exclusion information for ASD dataset. |
| Supplementary Table 4 | Detailed exclusion information for EOEC dataset. |
| Supplementary Table 5 | Demographic information of PD-off dataset. |
| Supplementary Table 6 | Demographic information of PD-on dataset. |
| Supplementary Table 7 | Demographic information of ASD dataset. |
| Supplementary Table 8 | Demographic information of MF dataset. |
| Supplementary Table 9 | Demographic information of EOEC dataset. |
| Supplementary Table 10 | SFC seeds. |
| Supplementary Figure 1 | Intersection mask of all subjects' coverage for between-group design dataset. |
| Supplementary Figure 2 | Intersection mask of all subjects' coverage for within-group design dataset. |
| Supplementary Figure 3 | Meta-analytic results of between-group comparison. |
| Supplementary Figure 4 | Meta-analytic results of within-group comparison. |
| Supplementary Figure 5 | Effect size of meta-analysis. |
| Supplementary Figure 6 | FNR, accuracy, and FDR for PD-off dataset. |
| Supplementary Figure 7 | FNR, accuracy, and FDR for PD-on dataset. |
| Supplementary Figure 8 | FNR, accuracy, and FDR for ASD dataset. |
| Supplementary Figure 9 | FNR, accuracy, and FDR for MF dataset. |
| Supplementary Figure 10 | FNR, accuracy, and FDR for EOEC dataset. |

### Supplementary Note 1. Information of each dataset

There were 5 datasets, including PD-off, PD-on, ASD, MF, and EOEC. The former 4 datasets were between-group design, and the EOEC was within-group design.

PD-off and PD-on datasets were from 20 research centers. Part of the datasets have been used in previous publications[1-24]. For PD-off, the levodopa was withdrawn for at least 12h before scanning. PD-off dataset consisted of 15 studies from 13 research institutes. PD-on dataset consisted of 9 studies from 9 institutes. Because of ethical issues, the raw data of most studies of PD-off dataset and PD-on dataset was not sent to Hangzhou Normal University. Instead, the data was analyzed in each research center by using the pipeline (rest_metabatch.m). The amplitude of low frequency fluctuation (ALFF), regional homogeneity (ReHo), and degree centrality (DC) images were sent to Hangzhou Normal University and were analyzed together.

ASD dataset was from the Autism Brain Imaging Data Exchange (ABIDE). The scanning parameters can be found in the ABIDE website (http://fcon_1000.projects.nitrc.org/indi/abide). The ABIDE initiative includes two large-scale collections: ABIDE Ⅰ and ABIDE Ⅱ. We included 27 studies from ABIDE Ⅰ and ABIDE Ⅱ.

MF (males vs. females) dataset consisted of 3 sub-datasets including MF_BNU (The Beijing_Zang from 1000 Functional Connectomes' Project), MF_CAMB (The Cambridg_Buckner from 1000 Functional Connectomes' Project), and MF_HZNU. Scan parameters of MF_BNU and MF_CAMB can be found in the FCP website (http://fcon_1000.projects.nitrc.org/fcpClassic/FcpTable.html). MF_HZNU sub-dataset was acquired using a GE Discovery MR-750 3.0 T scanner at the Center for Cognition and Brain Disorders of Hangzhou Normal University. The BOLD images were acquired using a gradient echo EPI pulse sequence with the following parameters: TR = 2000 ms, TE = 30ms, FOV = 220 × 220 $mm^2$, matrix = 64 × 64, flip angle = 90º, 43 axial slices, thickness/gap = 3.2/0 mm. The duration of resting-state scan was 8 minutes, and its includes 240 timepoints. From the 3 sub-datasets, we selected 360 subjects to generate 6 MF studies.

EOEC was eyes closed vs. eyes open within-group design. EOEC dataset consisted of 3 studies including EOEC_BNU[25], EOEC_HZNU01[26], and EOEC_HZNU02. The scanning parameters of EOEC_BNU and EOEC_HZNU01 can be found in the published papers[25,26]. EOEC_HZNU02 was collected using a GE Discovery MR-750 3.0 T scanner at the Center for Cognition and Brain Disorders of Hangzhou Normal University. The BOLD images were acquired using the same parameters as those of MF_HZNU. After subject exclusion, 107 subjects were included (EOEC_BNU: 43; EOEC_HNZU01: 33; EOEC_HNZU02: 31).

### Supplementary Note 2. ACKNOWLEDGMENTS

This work was supported by National Natural Science Foundation of China (81701671), Nanjing science and technology development projects (2014sc51701), Foundation of Natural Science of China (81471654), General Program of National Natural Science Foundation of China (61571047), the BRC for Mental Health at South London and Maudsley NHS Foundation Trust and by the Sackler Institute, Henry G Leong Endowed Professorship in Neurology, Henry G Leong Endowed Professorship in Neurology, National Institutes of Health (2R01AG006457, FBH), National Key R&D Program of China (2017YFC1309902), the National Natural Science Foundation of China (81671774 and 81630031), the Hundred Talents Program of the Chinese Academy of Sciences, and Beijing Municipal Science & Technology Commission (Z161100000216152), National Key Research and Development Plan (2016YFC1306600), National Natural Science Foundation of China (81771820, 81371519 and 81571654), National Nature Science Foundation of China(81701664), the Technology Innovation Program in Southwest Hospital (SWH2016JCYB-30), Natural Science Foundation of China (81520108016, 81661148045, and 31471084), "Qian Jiang Distinguished Professor" program, Natural Science Foundation of China (81571228), The National Key Research and Development Program of China (2016YFC1306503), Beijing Municipal Administration of Hospitals' Mission Plan (SML20150803), Beijing Municipal Science & Technology Commission (Z161100005116011, Z171100000117013), Beijing Municipal commission of Health and Family Planning (PXM2017_026283_000002).

**Supplementary Table 1. Detailed exclusion information for PD-off dataset.**

| | | Total n before exclusion | | Number of subjects excluded for detailed reasons | | | | | | | | Total n after exclusion | |
|---|---|---|---|---|---|---|---|---|---|---|---|---|---|
| | | | | Head motion | | Bad norm | | Small coverage | | Age or sex | | | |
| Institution | Study ID | PD | HC | PD | HC | PD | HC | PD | HC | PD | HC | PD | HC |
| BNU | Study_01 | 36 | 22 | 2 | 0 | 0 | 0 | 1 | 0 | 12 | 1 | 21 | 21 |
| BCU | Study_02 | 27 | 13 | 1 | 0 | 0 | 0 | 0 | 0 | 9 | 0 | 17 | 13 |
| TMMU | Study_03 | 50 | 26 | 0 | 0 | 0 | 0 | 0 | 0 | 12 | 0 | 38 | 26 |
| CMU | Study_04 | 34 | 40 | 0 | 0 | 0 | 0 | 0 | 0 | 0 | 5 | 34 | 35 |
| CCMU | Study_05 | 61 | 31 | 3 | 1 | 0 | 0 | 1 | 0 | 3 | 0 | 54 | 30 |
| | Study_06 | 17 | 24 | 0 | 1 | 0 | 0 | 0 | 0 | 0 | 0 | 17 | 23 |
| GGH | Study_07 | 40 | 27 | 0 | 0 | 0 | 0 | 0 | 0 | 4 | 0 | 36 | 27 |
| GPHCM | Study_08 | 16 | 20 | 0 | 0 | 0 | 0 | 0 | 0 | 0 | 0 | 16 | 20 |
| SCU | Study_09 | 17 | 20 | 0 | 0 | 0 | 1 | 0 | 0 | 0 | 0 | 17 | 19 |
| CNR | Study_10 | 12 | 15 | 0 | 0 | 0 | 0 | 0 | 0 | 1 | 4 | 11 | 11 |
| OHSU | Study_11 | 26 | 15 | 3 | 0 | 0 | 0 | 0 | 0 | 12 | 5 | 11 | 10 |
| UF | Study_12 | 39 | 20 | 1 | 0 | 0 | 0 | 6 | 1 | 1 | 0 | 31 | 19 |
| | Study_13 | 55 | 40 | 3 | 2 | 0 | 0 | 22 | 10 | 4 | 2 | 26 | 26 |
| SUN | Study_14 | 20 | 18 | 1 | 0 | 1 | 1 | 5 | 3 | 0 | 0 | 13 | 14 |
| SAHZU | Study_15 | 34 | 17 | 0 | 0 | 0 | 0 | 0 | 0 | 0 | 0 | 34 | 17 |
| 13 | 15 | 484 | 348 | 14 | 4 | 1 | 2 | 35 | 14 | 58 | 17 | 376 | 311 |

Some subjects of PD-off dataset were excluded for the following reasons. 1) Head motion bigger than 3 mm or 3 degrees. 2) Bad spatial normalization. 3) Scans covered smaller than 97% of the whole brain. To avoid deleting too many subjects, a coverage of 97% was used. 4) The age or sex was not matched. Some subjects were excluded to get the comparison groups matched for age ($P > 0.5$, Two-sample t-test) and sex ($P > 0.5$, Chi-square test) in each study of PD-off dataset. Head motion, the exclusion criteria were bigger than 3 mm or 3 degrees. Bad norm, bad spatial normalization. Small coverage, exclusion criteria were smaller than 97% of whole brain. Sex or age, the sex or age was not matched. Some studies from PD-off dataset didn't have handedness information and hence no subject was excluded for handedness. PD-off, Parkinson's disease off levodopa. HC, healthy controls. BNU, State Key Laboratory of Cognitive Neuroscience and Learning, Beijing Normal University. BCU, Department of Medicine (Neurology) and Pacific Parkinson's Research Centre, University of British Columbia. TMMU, Department of Radiology, Southwest Hospital, Third Military Medical University, Chongqing. CMU, Department of Radiology, The First Affiliated Hospital of China Medical University. CCMU, Beijing Institute of Geriatrics, Xuanwu Hospital, Capital Medical University. GGH, Department of Radiology, Guangdong Academy of Medical Sciences, Guangdong General Hospital. GPHCM, Department of Radiology, Guangdong Provincial Hospital of Chinese Medicine. SCU, Department of Neurology, West China Hospital, SiChuan University. CNR, Consiglio Nazionale delle Ricerche of Catanzaro Italy, IBFM. OHSU, Department of Neurology, School of Medicine, Oregon Health & Science University. UF, Department of Applied Physiology and Kinesiology, University of Florida. SUN, Department of Neurology, Second University of Naples. SAHZU, The Second Affiliated Hospital of Zhejiang University School of Medicine.

### Supplementary Table 2. Detailed exclusion information for PD-on dataset.

| | | Total n before exclusion | | Number of subjects excluded for detailed reasons | | | | | | | | Total n after exclusion | |
|---|---|---|---|---|---|---|---|---|---|---|---|---|---|
| | | | | Head motion | | Bad norm | | Small coverage | | Age or sex | | | |
| Institution | Study ID | PD | HC | PD | HC | PD | HC | PD | HC | PD | HC | PD | HC |
| BCU | Study_01 | 52 | 13 | 2 | 0 | 0 | 0 | 0 | 0 | 22 | 0 | 28 | 13 |
| US | Study_02 | 39 | 11 | 1 | 0 | 0 | 0 | 11 | 4 | 9 | 0 | 18 | 7 |
| NDFU | Study_03 | 36 | 16 | 4 | 1 | 0 | 0 | 18 | 6 | 4 | 0 | 10 | 9 |
| UU | Study_04 | 12 | 12 | 0 | 0 | 0 | 0 | 0 | 1 | 0 | 0 | 12 | 11 |
| CEIT | Study_05* | 32 | 18 | 9 | 1 | 0 | 0 | 23 | 15 | 0 | 0 | 0 | 2 |
| CNR | Study_06 | 11 | 15 | 2 | 0 | 0 | 0 | 0 | 0 | 0 | 3 | 9 | 12 |
| HKU | Study_07 | 44 | 25 | 8 | 1 | 0 | 0 | 0 | 0 | 9 | 0 | 27 | 24 |
| UW | Study_08 | 24 | 21 | 0 | 2 | 1 | 1 | 5 | 2 | 2 | 1 | 16 | 15 |
| NMU | Study_09 | 20 | 20 | 1 | 1 | 0 | 0 | 4 | 2 | 0 | 3 | 15 | 14 |
| SUN | Study_10 | 20 | 18 | 0 | 0 | 0 | 1 | 8 | 3 | 1 | 4 | 11 | 10 |
| 10 | 10 | 290 | 169 | 27 | 6 | 1 | 2 | 69 | 33 | 47 | 11 | 146 | 117 |

Some subjects of PD-on dataset were excluded for the following reasons. 1) Head motion bigger than 3 mm or 3 degrees. 2) Bad spatial normalization. 3) Scans covered smaller than 97% of the whole brain. To avoid deleting too many subjects, a coverage of 97% was used. 4) The age or sex was not matched. Some subjects were excluded to get the comparison groups matched for age (*P* > 0.5, Two-sample t-test) and sex (*P* > 0.5, Chi-square test) in each study of PD-on dataset. Head motion, the exclusion criteria were bigger than 3 mm or 3 degrees. Bad norm, bad spatial normalization. Small coverage, exclusion criteria were smaller than 97% of whole brain. Sex or age, the sex or age was not matched. Some studies from PD-on dataset didn't have handedness information and hence no subject was excluded for handedness. coverage. PD-on, Parkinson's disease on levodopa. HC, healthy controls. BCU, Department of Medicine (Neurology) and Pacific Parkinson's Research Centre, University of British Columbia. US, Brain and Mind Research Institute, The University of Sydney. NDFU, Université Lille Nord de France. UU, Department of Neurology, University of Ulm. CNR, Consiglio Nazionale delle Ricerche of Catanzaro Italy, IBFM. HKU, Division of Neurology, Department of Medicine, Queen Mary Hospital, The University of Hong Kong. UW, Department of Radiology, University of Washington. NMU, Department of Neurology, Affiliated Brain Hospital of Nanjing Medical University. SUN, The Second Affiliated Hospital of Zhejiang University School of Medicine.

*: Study_05 was excluded because no enough participants were left due to small scanning

### Supplementary Table 3. Detailed exclusion information for ASD dataset

| | | Total n before exclusion | | Number of subjects excluded for detailed reasons | | | | | | | | | | | | | | Total n after exclusion | |
|---|---|---|---|---|---|---|---|---|---|---|---|---|---|---|---|---|---|---|---|
| | | | | Not right | | Scan para | | Head motion | | Bad Norm | | Small coverage | | Age or sex | | Other | | | |
| Institution name | Study ID | ASD | HC | ASD | HC | ASD | HC | ASD | HC | ASD | HC | ASD | HC | ASD | HC | ASD | HC | ASD | HC |
| ABIDE Ⅰ CALTECH | Study_01 | 19 | 19 | 5 | 4 | 0 | 0 | 0 | 1 | 0 | 0 | 6 | 1 | 0 | 2 | 0 | 0 | 8 | 11 |
| ABIDE Ⅰ KKI | Study_02 | 22 | 33 | 4 | 5 | 4 | 0 | 4 | 7 | 0 | 0 | 0 | 0 | 0 | 5 | 0 | 0 | 10 | 16 |
| ABIDE Ⅰ LEUVEN | Study_03 | 29 | 35 | 3 | 5 | 0 | 0 | 1 | 2 | 0 | 0 | 10 | 7 | 0 | 0 | 0 | 0 | 15 | 21 |
| ABIDE Ⅰ NYU | Study_04 | 79 | 105 | 18 | 6 | 0 | 0 | 4 | 7 | 1 | 0 | 3 | 8 | 0 | 12 | 4 | 1 | 49 | 71 |
| ABIDE Ⅰ OHSU | Study_05 | 13 | 15 | 1 | 0 | 0 | 0 | 0 | 2 | 0 | 0 | 0 | 0 | 1 | 2 | 1 | 1 | 10 | 10 |
| ABIDE Ⅰ OLIN | Study_06 | 20 | 16 | 4 | 2 | 0 | 0 | 4 | 5 | 0 | 0 | 1 | 1 | 1 | 0 | 0 | 0 | 10 | 8 |
| ABIDE Ⅰ SDSU | Study_07 | 14 | 22 | 1 | 3 | 0 | 0 | 2 | 1 | 0 | 0 | 0 | 0 | 0 | 8 | 0 | 0 | 11 | 10 |
| ABIDE Ⅰ STANFORD | Study_08‡ | 20 | 20 | 5 | 2 | 0 | 0 | 1 | 4 | 0 | 0 | 7 | 5 | 0 | 1 | 0 | 0 | 7 | 8 |
| ABIDE Ⅰ TRINITY | Study_09 | 24 | 25 | 0 | 0 | 0 | 0 | 4 | 2 | 0 | 0 | 0 | 0 | 0 | 0 | 1 | 0 | 19 | 23 |
| ABIDE Ⅰ UCLA | Study_10 | 62 | 47 | 6 | 4 | 0 | 0 | 9 | 4 | 5 | 8 | 2 | 1 | 0 | 0 | 0 | 1 | 40 | 29 |
| ABIDE Ⅰ UM_2 | Study_11 | 13 | 22 | 2 | 2 | 0 | 0 | 3 | 3 | 0 | 0 | 1 | 8 | 1 | 3 | 0 | 0 | 6 | 6 |
| ABIDE Ⅰ USM | Study_12 | 58 | 43 | 7 | 2 | 2 | 0 | 18 | 13 | 1 | 0 | 5 | 4 | 0 | 0 | 0 | 8 | 25 | 16 |
| ABIDE Ⅰ YALE | Study_13 | 28 | 28 | 6 | 4 | 0 | 0 | 3 | 0 | 7 | 12 | 2 | 1 | 0 | 0 | 0 | 0 | 10 | 11 |
| ABIDE Ⅱ BNI_1 | Study_14 | 29 | 29 | 0 | 0 | 2 | 0 | 2 | 7 | 6 | 7 | 0 | 0 | 1 | 0 | 2 | 0 | 16 | 15 |
| ABIDE Ⅱ EMC_1 | Study_15 | 27 | 27 | 5 | 6 | 0 | 0 | 8 | 7 | 0 | 0 | 0 | 0 | 1 | 1 | 0 | 0 | 13 | 13 |
| ABIDE Ⅱ ETH_1 | Study_16 | 13 | 24 | 0 | 0 | 0 | 0 | 5 | 3 | 0 | 0 | 0 | 1 | 0 | 5 | 0 | 0 | 8 | 15 |
| ABIDE Ⅱ GU_1 | Study_17 | 51 | 55 | 8 | 3 | 0 | 0 | 15 | 11 | 8 | 13 | 6 | 3 | 0 | 11 | 0 | 1 | 14 | 13 |
| ABIDE Ⅱ IP_1 | Study_18 | 22 | 34 | 1 | 7 | 0 | 0 | 1 | 2 | 0 | 0 | 4 | 5 | 2 | 7 | 0 | 0 | 14 | 13 |
| ABIDE Ⅱ IU_1 | Study_19 | 20 | 20 | 5 | 3 | 0 | 0 | 0 | 0 | 0 | 0 | 0 | 0 | 0 | 0 | 0 | 0 | 15 | 17 |
| ABIDE Ⅱ KKI_1 | Study_20 | 56 | 155 | 10 | 22 | 0 | 9 | 16 | 33 | 1 | 4 | 0 | 2 | 0 | 11 | 0 | 0 | 29 | 74 |

| | | | | | | | | | | | | | | | | | | | |
|---|---|---|---|---|---|---|---|---|---|---|---|---|---|---|---|---|---|---|---|
| ABIDE Ⅱ NYU_1 | Study_21 | 48 | 30 | 21 | 2 | 1 | 0 | 0 | 1 | 2 | 3 | 1 | 1 | 0 | 0 | 0 | 0 | 23 | 23 |
| ABIDE Ⅱ OHSU_1 | Study_22 | 37 | 56 | 2 | 1 | 0 | 0 | 2 | 2 | 1 | 0 | 1 | 0 | 3 | 26 | 0 | 0 | 28 | 27 |
| ABIDE Ⅱ OILH_2 | Study_23 | 24 | 35 | 8 | 3 | 0 | 1 | 2 | 2 | 0 | 1 | 0 | 0 | 2 | 16 | 2 | 3 | 10 | 9 |
| ABIDE Ⅱ SDSU_1 | Study_24 | 33 | 25 | 6 | 5 | 0 | 0 | 1 | 0 | 0 | 0 | 0 | 0 | 2 | 0 | 0 | 0 | 24 | 20 |
| ABIDE Ⅱ TCD_1 | Study_25 | 21 | 21 | 0 | 0 | 1 | 0 | 6 | 3 | 0 | 0 | 1 | 1 | 1 | 5 | 0 | 0 | 12 | 12 |
| ABIDE Ⅱ UCD_1 | Study_26 | 18 | 14 | 1 | 0 | 0 | 0 | 1 | 0 | 0 | 0 | 4 | 2 | 0 | 0 | 0 | 0 | 12 | 12 |
| ABIDE Ⅱ UCLA_1 | Study_27 | 16 | 16 | 2 | 2 | 0 | 0 | 1 | 1 | 1 | 0 | 0 | 0 | 3 | 5 | 0 | 0 | 9 | 8 |
| | 27 | 816 | 971 | 131 | 93 | 10 | 10 | 113 | 123 | 33 | 48 | 54 | 51 | 18 | 120 | 10 | 15 | 447 | 511 |

ASD dataset was from ABIDE (http://fcon_1000.projects.nitrc.org/indi/abide). ABIDE Ⅰ and Ⅱ involved 17 and 19 sites, respectively. ABIDE Ⅱ, Stanford University and ABIDE Ⅱ, University of Miami were not included because we failed to download these data. Some studies were excluded for the following reasons. 1) ABIDE Ⅰ, Carnegie Mellon University. Too many subjects were excluded due to realignment error or segment error. 2) ABIDE Ⅰ, University of Michigan, Sample 1. Too many subjects were excluded due to realignment error or segment error. 3) ABIDE Ⅰ, Social Brain Lab BCN NIC UMC Groningen and Netherlands Institute for Neurosciences. Only one subject was left after small coverage exclusion. 4) ABIDE Ⅰ, Ludwig Maximilians University Munich (ABIDE Ⅰ MaxMun). We divided ABIDE Ⅰ MaxMun into two studies because two sets of parameters were used (slice number 28 and 40). After small coverage exclusion, only 2 and 10 participants were left for the two sub-datasets of ABIDE1-MaxMun (sub-dataset 1 slice number = 28, and sub-dataset 2 slice number = 40). 5) ABIDE Ⅰ University of Pittsburgh School of Medicine. Only 11 subjects were left after age and sex matched. 6) ABIDE Ⅱ University of Utah School of Medicine. Only 7 subjects were left after age and sex were matched. 7) ABIDE Ⅱ Katholieke Universiteit Leuven. No control groups. 8) ABIDE Ⅱ University of Pittsburgh School of Medicine, Longitudinal Sample. Only 2 subjects were left after small coverage exclusion. 9) ABIDE Ⅱ NYU Langone Medical Center, Sample 2. No control groups. 10) ABIDE Ⅱ University of California Los Angeles, Longitudinal Sample. Only 6 subjects were left after age and sex were matched. Some subjects of ASD dataset were excluded for the following reasons. 1) Non-right handedness subjects. 2) Scan parameter (slice number, timepoints) was different from most subjects within a study. 3) Head motion bigger than 3 mm or 3 degrees. 4) Bad spatial normalization. 5) Scans covered smaller than 97% of the whole brain. To avoid deleting too many subjects, a coverage of 97% was used. 6) The age or sex was not matched. Some subjects were excluded to get the comparison groups matched for age ($P > 0.5$, Two-sample t-test) and sex ($P > 0.5$, Chi-square test) in each study of ASD dataset. Not right, non-right handedness subjects. Head motion, the exclusion criteria were bigger than 3 mm or 3 degrees. Bad norm, bad spatial normalization. Small coverage, exclusion criteria were smaller than 97% of whole brain. Sex or age, the sex or age was not matched. Other, these subjects were excluded due to other reasons as listed below. 1) Study_04, four subjects were excluded due to realignment error (Subject number: 50953, 50975, 50980 and 51108). One subject was excluded due to “new segment” error (Subject number: 50992). 2) Study_05, two subjects did not have resting-state fMRI data (Subject number: 50165 and 50155). 3) Study_09, one subject was excluded due to realignment error (Subject number: 50238). 4) Study_10, one subject was excluded due to realignment error (Subject number: 51259). 5) Study_12, eight subjects did not have resting-state fMRI data (Subject number: 50452, 50457, 50458, 50459, 50460, 50461, 50462, and 50465). 6) Study_14, two subjects were excluded due to realignment error (Subject number: 29006 and 29007). 7) Study_17, one subject was excluded due to new segment error (Subject number: 28781). 8) Study_23, one subject did not have structural MRI data (Subject number: 28682). Six subjects did not have resting-state fMRI data (Subject number: 28681, 28683, 28687, 28711, 28712 and 28713). ASD, autism spectrum disorder. HC, healthy controls. ABIDE, Autism Brain Imaging Data Exchange. CALTECH, California Institute of Technology. KKI, Kennedy Krieger Institute. LEUVEN, University of Leuven. NYU, NYU Langone Medical Center. OHSU, Oregon Health and Science University. OLIN, Olin, Institute of Living at Hartford Hospital. SDSU, San Diego State University. STANFORD, Stanford University. TRINITY, Trinity Centre for Health Sciences. UCLA, University of California, Los Angeles. UM_2, University of Michigan, Sample 2. USM, University of Utah School of Medicine. YALE, Yale Child Study Center. BNI_1, Barrow Neurological Institute. EMC_1, Erasmus University Medical Center Rotterdam. ETH_1, ETH Zürich. GU_1, Georgetown University. IP_1, Institute Pasteur and Robert Debré Hospital. IU_1, Indiana University. KKI_1, Kennedy Krieger Institute. NYU_1, NYU Langone Medical Center, Sample 1. OHSU_1, Oregon Health and Science University. OILH_2, Olin Neuropsychiatry Research Center, Institute of Living at Hartford Hospital. SDSU_1, San Diego State University. TCD_1, Trinity Centre for Health Sciences. UCD_1, University of California Davis. UCLA_1, University of California Los Angeles.

‡For Study08, the timepoints were not same for all subjects (Twenty subjects had 240 timepoints. One subject had 181 timepoints. Two subjects had 238 timepoints. Seventeen subjects had 180 timepoints). We used 180 timepoints for all subjects.

**Supplementary Table 4. Detailed exclusion information for EOEC dataset**.

| | Total n before exclusion | Number of subjects excluded for detailed reasons | | | | Total n after exclusion |
|---|---|---|---|---|---|---|
| | | Not right | Head motion | Bad norm | Small coverage | |
| EOEC_BNU | 47 | 3 | 0 | 0 | 1 | 43 |
| EOEC_HZNU01 | 34 | 0 | 0 | 1 | 0 | 33 |
| EOEC_HZNU02 | 31 | 0 | 0 | 0 | 0 | 31 |

Some subjects of EOEC dataset were excluded for the following reasons. 1) Non-right handedness subjects. 2) Head motion bigger than 3 mm or 3 degrees. 3) Bad spatial normalization. 4) Scans covered smaller than 97% of the whole brain. To avoid deleting too many subjects, a coverage of 97% was used. The age and sex were already matched for all the studies of EOEC dataset. Not right, non-right handedness subjects. Head motion, the exclusion criteria were bigger than 3 mm or 3 degrees. Bad norm, bad spatial normalization. Small coverage, exclusion criteria were smaller than 97% of whole brain. EOEC, eye open vs. eyes closed.

### Supplementary Table 5. Demographic information of PD-off dataset.

| Study ID | PD | | | HC | | | Sex *P* | Age *P* |
|---|---|---|---|---|---|---|---|---|
| | Male:female | Right:non-right | Age | Male:female | Right:non-right | Age | | |
| Study_01 | 12:9 | 21:0 | 57.3±8.2 | 13:8 | 21:0 | 56.2±6.4 | 0.75 | 0.65 |
| Study_02 | 11:6 | 16:1 | 58.9±8.9 | 7:6 | 13:0 | 58.0±6.0 | 0.55 | 0.75 |
| Study_03 | 16:22 | 38:0 | 58.8±11.5 | 9:17 | 26:0 | 58.8±7.8 | 0.55 | 1.00 |
| Study_04 | 18:16 | 34:0 | 61.8±10.0 | 16:19 | 35:0 | 62.1±7.8 | 0.55 | 0.88 |
| Study_05 | 29:25 | 54:0 | 64.3±7.4 | 15:15 | \ | 65.2±6.5 | 0.74 | 0.55 |
| Study_06 | 6:11 | 16:1 | 60.4±9.2 | 10:13 | 22:1 | 59.1±9.4 | 0.60 | 0.67 |
| Study_07 | 15:21 | 36:0 | 62.4±8.5 | 11:16 | 27:0 | 63.8±9.9 | 0.94 | 0.56 |
| Study_08 | 9:7 | 16:0 | 60.5±11.8 | 11:9 | \ | 59.2±8.7 | 0.94 | 0.71 |
| Study_09 | 6:11 | 17:0 | 64.0±5.0 | 7:12 | 19:0 | 63.7±7.0 | 0.92 | 0.88 |
| Study_10 | 7:4 | 11:0 | 66.0±6.6 | 7:4 | 11:0 | 64.2±6.0 | 1.00 | 0.51 |
| Study_11 | 7:4 | \ | 67.4±8.6 | 5:5 | \ | 67.5±8.8 | 0.53 | 0.97 |
| Study_12 | 24:7 | 29:2 | 60.5±10.5 | 15:4 | 17:2 | 62.2±9.7 | 0.90 | 0.56 |
| Study_13 | 10:16 | 23:3 | 61.5±9.6 | 8:18 | 23:3 | 61.5±10.1 | 0.56 | 0.98 |
| Study_14 | 4:9 | \ | 62.2±8.4 | 6:8 | \ | 61.4±7.1 | 0.52 | 0.79 |
| Study_15 | 17:17 | \ | 59.2±9.3 | 9:8 | \ | 59.2±9.9 | 0.84 | 0.99 |

Totally 15 studies of PD-off were included in the meta-analysis of WBVW metrics. Due to ethnical issues, SFC was not performed for PD-off. Some studies from PD-off dataset didn't have handedness information and hence no subject was excluded for handedness.

"\", no handedness information. PD-off, Parkinson's disease off levodopa. HC, healthy controls.

**Supplementary Table 6. Demographic information of PD-on dataset.**

| Study ID | PD | | | HC | | | Sex *P* | Age *P* |
|---|---|---|---|---|---|---|---|---|
| | Male:female | Right:non-right | Age | Male:female | Right:non-right | Age | | |
| Study_01 | 18:10 | 26:2 | 59.6±8.4 | 7:6 | 13:0 | 58.0±6.0 | 0.52 | 0.53 |
| Study_02 | 10:8 | 15:3 | 69.7±7.1 | 3:4 | \ | 69.7±8.4 | 0.57 | 0.99 |
| Study_03 | 8:2 | 10:0 | 55.6±9.9 | 8:1 | 9:0 | 54.9±8.3 | 0.60 | 0.87 |
| Study_04 | 6:6 | 12:0 | 66.4±6.4 | 5:6 | 11:0 | 66.1±5.7 | 0.83 | 0.93 |
| Study_06 | 6:3 | 9:0 | 65.7±7.0 | 8:4 | 12:0 | 63.8±5.9 | 1.00 | 0.50 |
| Study_07 | 16:11 | \ | 61.7±6.9 | 12:12 | \ | 61.5±5.0 | 0.51 | 0.92 |
| Study_08 | 9:7 | 12:4 | 61.8±8.1 | 7:8 | 13:2 | 61.3±8.9 | 0.59 | 0.88 |
| Study_09 | 3:12 | \ | 58.8±7.7 | 4:10 | \ | 58.9±5.0 | 0.59 | 0.96 |
| Study_10 | 7:4 | \ | 64.0±6.5 | 5:5 | \ | 64.2±6.0 | 0.53 | 0.94 |

Totally 10 studies of PD-on were included in the meta-analysis of WBVW metrics. Due to ethnical issues, PD-on was not included in meta-analysis for SFC. Some studies from PD-on dataset didn't have handedness information and hence no subject was excluded for handedness.

"\", no handedness information. PD-on, Parkinson's disease on levodopa. HC, healthy controls.

**Supplementary Table 7. Demographic information of ASD dataset.**

| Study ID | ASD | | HC | | Sex *P* | Age *P* |
|---|---|---|---|---|---|---|
| | Male:female | Age | Male:female | Age | | |
| Study_01 | 7:1 | 28.0±12.4 | 9:2 | 30.0±12.7 | 0.74 | 0.74 |
| Study_02 | 9:1 | 9.8±1.7 | 13:3 | 10.2±1.1 | 0.55 | 0.55 |
| Study_03 | 14:1 | 19.8±4.4 | 19:2 | 20.0±5.1 | 0.76 | 0.90 |
| Study_04 | 42:7 | 14.9±7.0 | 58:13 | 15.0±6.1 | 0.56 | 0.98 |
| Study_05 | 10:0 | 10.8±1.7 | 10:0 | 10.6±0.9 | 1.00 | 0.74 |
| Study_06 | 8:2 | 15.9±3.0 | 6:2 | 15.5±2.9 | 0.80 | 0.78 |
| Study_07 | 11:0 | 15.2±1.7 | 10:0 | 15.0±1.1 | 1.00 | 0.75 |
| Study_08 | 6:1 | 10.0±1.9 | 6:2 | 10.0±1.5 | 0.60 | 0.99 |
| Study_09 | 19:0 | 17.0±2.7 | 23:0 | 17.5±3.7 | 1.00 | 0.60 |
| Study_10 | 35:5 | 13.2±2.3 | 24:5 | 13.2±1.9 | 0.58 | 0.96 |
| Study_11 | 6:0 | 14.8±1.6 | 6:0 | 15.1±0.7 | 1.00 | 0.68 |
| Study_12 | 25:0 | 23.3±7.7 | 16:0 | 23.4±8.3 | 1.00 | 0.97 |
| Study_13 | 6:4 | 14.0±2.4 | 7:4 | 13.7±2.2 | 0.86 | 0.76 |
| Study_14 | 16:0 | 33.9±15.4 | 15:0 | 37.7±15.9 | 1.00 | 0.51 |
| Study_15 | 10:3 | 8.2±0.9 | 10:3 | 8.1±0.7 | 1.00 | 0.74 |
| Study_16 | 8:0 | 20.8±4.3 | 15:0 | 22.0±4.0 | 1.00 | 0.51 |
| Study_17 | 12:2 | 10.9±1.7 | 10:3 | 10.9±2.1 | 0.56 | 1.00 |
| Study_18 | 7:7 | 15.7±4.9 | 5:8 | 16.6±6.3 | 0.55 | 0.68 |
| Study_19 | 12:3 | 24.9±10.8 | 13:4 | 23.5±5.0 | 0.81 | 0.63 |
| Study_20 | 20:9 | 10.5±1.5 | 46:28 | 10.5±1.1 | 0.52 | 1.00 |
| Study_21 | 21:2 | 10.2±6.0 | 22:1 | 10.1±3.5 | 0.55 | 0.94 |
| Study_22 | 22:6 | 11.6±2.1 | 21:6 | 11.3±1.6 | 0.94 | 0.59 |
| Study_23 | 10:0 | 21.9±3.7 | 9:0 | 22.4±2.7 | 1.00 | 0.72 |
| Study_24 | 20:4 | 13.1±3.4 | 18:2 | 13.1±2.9 | 0.52 | 0.99 |
| Study_25 | 12:0 | 14.6±3.4 | 12:0 | 15.2±2.3 | 1.00 | 0.62 |
| Study_26 | 9:3 | 14.9±1.9 | 9:3 | 14.7±1.7 | 1.00 | 0.77 |
| Study_27 | 8:1 | 11.0±1.7 | 7:1 | 10.5±2.1 | 0.93 | 0.60 |

### Supplementary Table 8. Demographic information of MF dataset.

| Sub dataset | Study ID | Age for male | Age for female | *P* |
|---|---|---|---|---|
| MF_BNU | Study_01 | 20.8±2.0 | 20.8±2.0 | 0.95 |
| MF_CAMB | Study_02 | 20.8±1.4 | 21.0±1.4 | 0.52 |
| MF_CAMB | Study_03 | 21.0±1.7 | 20.7±1.7 | 0.60 |
| MF_HZNU | Study_04 | 30.8±9.2 | 29.6±9.5 | 0.64 |
| MF_HZNU | Study_05 | 32.5±9.8 | 31.4±11.5 | 0.69 |
| MF_HZNU | Study_06 | 30.3±11.4 | 32.6±10.8 | 0.44 |

The MF dataset was composed of 3 sub-datasets with large sample size of young adults. The current study aimed to investigate the false discovery rate of smaller sample size. Therefore, we extracted 6 studies, each containing 30 males and 30 females with the age matched. MF, healthy male vs. female.

**Supplementary Table 9. Demographic information of EOEC dataset.**

| Study name | Study ID | Age | Male:female |
|---|---|---|---|
| EOEC_BNU | Study_01 | 22.5±2.2 | 23:24 |
| EOEC_HZNU01 | Study_02 | 23.5±2.0 | 16:17 |
| EOEC_HZNU02 | Study_03 | 21.8±1.8 | 16:15 |

EOEC dataset was within-group design. Totally 3 studies of EOEC were included in the meta-analysis.

### Supplementary Table 10. SFC seeds.

| | Seed ID | Coordinates | | | Effect size | Region |
|---|---|---|---|---|---|---|
| | | x | y | z | | |
| ASD | Seed_01 | -24 | -20 | 30 | -0.35 | Undefined |
| | Seed_02 | 24 | -60 | -26 | 0.31 | Cerebelum_6_R |
| | Seed_03 | 12 | -96 | 12 | -0.31 | Cuneus_R |
| | Seed_04 | 30 | 48 | -4 | 0.30 | Frontal_Mid_Orb_R |
| | Seed_05 | -20 | 30 | 6 | -0.30 | Undefined |
| | Seed_06 | -18 | -96 | 6 | -0.29 | Occipital_Mid_L* |
| | Seed_07 | 30 | 14 | -36 | 0.29 | Temporal_Pole_Mid_R |
| | Seed_08 | -12 | 14 | -20 | 0.29 | Frontal_Sup_Orb_L |
| | Seed_09 | 36 | 30 | 48 | 0.28 | Frontal_Mid_R |
| | Seed_10 | 56 | -60 | -24 | -0.27 | Temporal_Inf_R |
| FM | Seed_01 | 0 | -46 | 14 | -0.70 | Undefined |
| | Seed_02 | -32 | 30 | -18 | 0.65 | Frontal_Inf_Orb_L |
| | Seed_03 | -54 | -62 | -40 | -0.63 | Cerebelum_Crus1_L |
| | Seed_04 | 26 | -88 | -28 | 0.60 | Cerebelum_Crus1_R |
| | Seed_05 | -22 | 68 | 0 | -0.60 | Frontal_Sup_L |
| | Seed_06 | 36 | 30 | -18 | 0.59 | Frontal_Inf_Orb_R |
| | Seed_07 | 18 | 0 | -12 | 0.55 | Undefined |
| | Seed_08 | 44 | -60 | 24 | -0.53 | Angular_R |
| | Seed_09 | 36 | 0 | 14 | 0.53 | Insula_R* |
| | Seed_10 | 12 | 6 | 64 | -0.52 | Supp_Motor_Area_R |
| EOEC | Seed_01 | 12 | -34 | 68 | -0.82 | Postcentral_R |
| | Seed_02 | 36 | -78 | 16 | 0.66 | Occipital_Mid_R* |
| | Seed_03 | 12 | -48 | 6 | -0.64 | Precuneus_R |
| | Seed_04 | -12 | -24 | -2 | -0.62 | Thalamus_L |
| | Seed_05 | -50 | -72 | 6 | -0.61 | Temporal_Mid_L |
| | Seed_06 | 18 | 60 | -4 | 0.61 | Frontal_Sup_Orb_R |
| | Seed_07 | -36 | -82 | 20 | 0.61 | Occipital_Mid_L |
| | Seed_08 | -10 | -48 | 4 | -0.59 | Calcarine_L |
| | Seed_09 | 44 | 6 | 24 | 0.56 | Frontal_Inf_Oper_R |
| | Seed_10 | 60 | -34 | 28 | 0.55 | SupraMarginal_R |

The selection of seed region of interest (ROI) for SFC analysis varies a lot in previous studies. We selected 10 seed ROIs for each dataset, which were from the 10 most abnormal ALFF clusters of meta-analytic results of each dataset. A spherical ROI (radius = 6mm, centered at the peak voxel of each cluster) was taken as the seed ROI for SFC. After preprocessing, Pearson correlation was calculated between the seed-ROI time course and the time course of each voxel in the whole brain. The Pearson correlation coefficient was Fisher *Z*-transformed.The label for brain regions was from automated anatomical labeling (AAL) [27] template by using a free software Xjview 8.14 (http://www.alivelearn.net/xjview).

* Representative SFC was shown in the main content of the article.

**Supplementary Figure 1. Intersection mask of all subjects' coverage for between-group design dataset.**

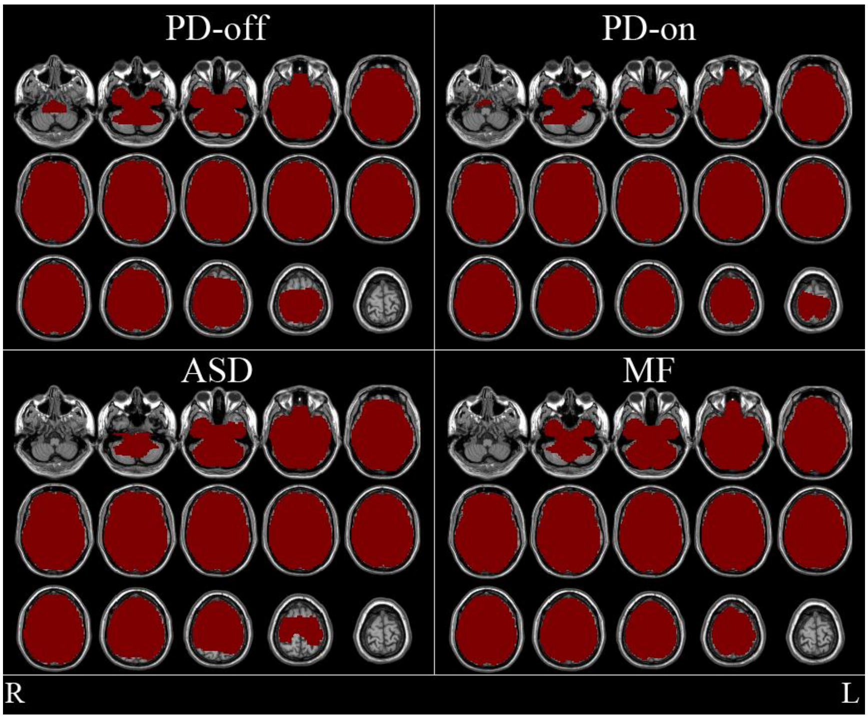


After spatial normalization, the mean EPI image across timepoints for each subject of each dataset of between-group design was calculated and saved as a binary image, i.e., the value outside the field of view was set zero. An intersection mask for each dataset was obtained by combining all subjects' binary images and a whole brain mask which was provided in the software DPABI V2.3[28]. All statistical analyses were performed within the dataset-specific intersection mask.

**Supplementary Figure 2. Intersection mask of all subjects' coverage for within-group design dataset.**

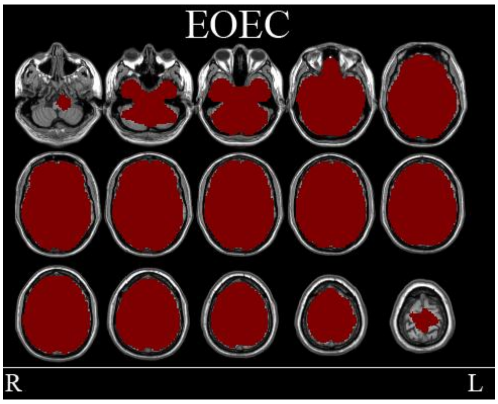


After spatial normalization, the mean EPI image across timepoints for each subject of each dataset of within-group design was calculated and saved as a binary image, i.e., the value outside the field of view was set zero. An intersection mask for each dataset was obtained by combining all subjects' binary images and a whole brain mask which was provided in the software DPABI V2.3[28]. All statistical analyses were performed within the dataset-specific intersection mask.

**Supplementary Figure 3. Meta-analytic results of between-group comparison.**

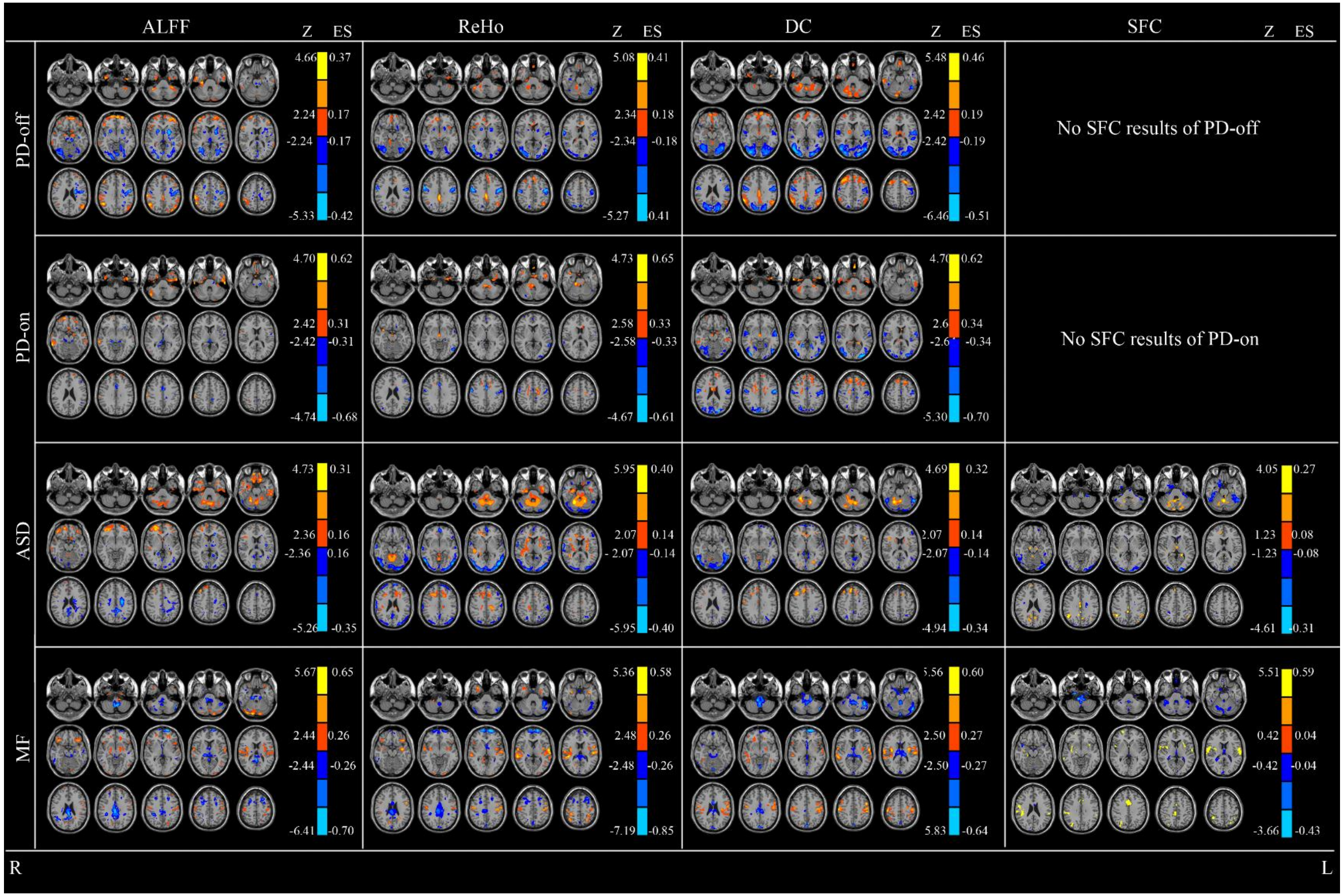

Warm colors indicate higher metrics (i.e. ALFF, ReHo, DC, and SFC) in PD than control (PD-off and PD-on), ASD than control, and males than females. Cold colors indicate the opposite. A combination threshold $P < 0.005$, $z > 1$, and cluster size > 10 voxels was used. The Z coordinates were from -47 to +64 with a step of 8 mm. ES, effect size (Hedges' $g$). L, left side of the brain. R, right side of the brain. PD-off, Parkinson's disease off levodopa vs. healthy controls (HC). PD-on, PD on levodopa vs. HC. ASD, autism spectrum disorder vs HC. MF, healthy male vs. female. ALFF, Amplitude of low frequency fluctuation. ReHo, Regional homogeneity. DC, Degree centrality. SFC, Seed-based functional connectivity. Because of ethical issues, SFC was not analyzed for PD-off and PD-on.

**Supplementary Figure 4. Meta-analytic results of within-group comparison.**

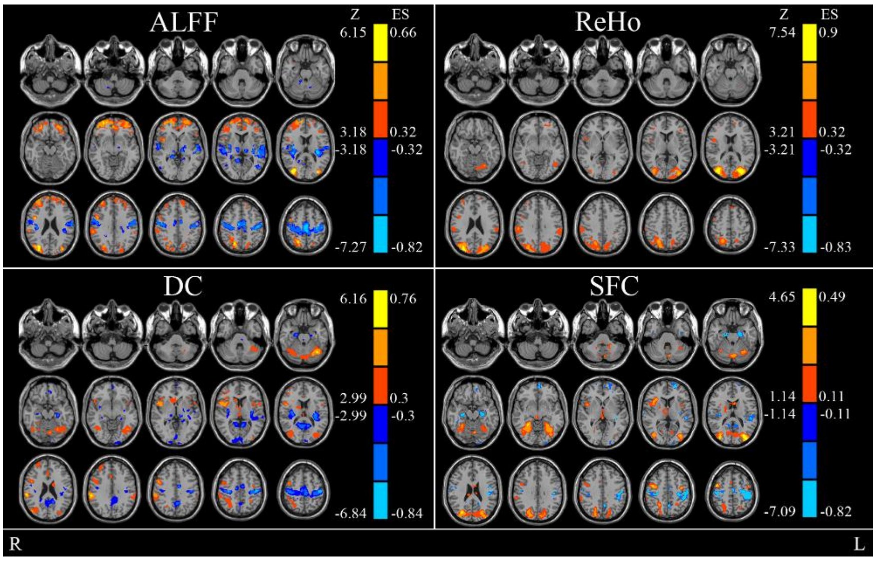


Warm colors indicate higher metrics (i.e., ALFF, ReHo, DC, and SFC) in eyes open than eyes closed. Cold colors indicate the opposite. A combination threshold $P < 0.005$, $z > 1$, and cluster size > 10 voxels was used. The Z coordinates were from -47 to +64 with a step of 8 mm. ES, effect size (Hedges' *g*). L, left side of the brain. R, right side of the brain. ALFF, Amplitude of low frequency fluctuation. ReHo, Regional homogeneity. DC, Degree centrality. SFC, Seed-based functional connectivity.

**Supplementary Figure 5. Effect size of meta-analysis.**

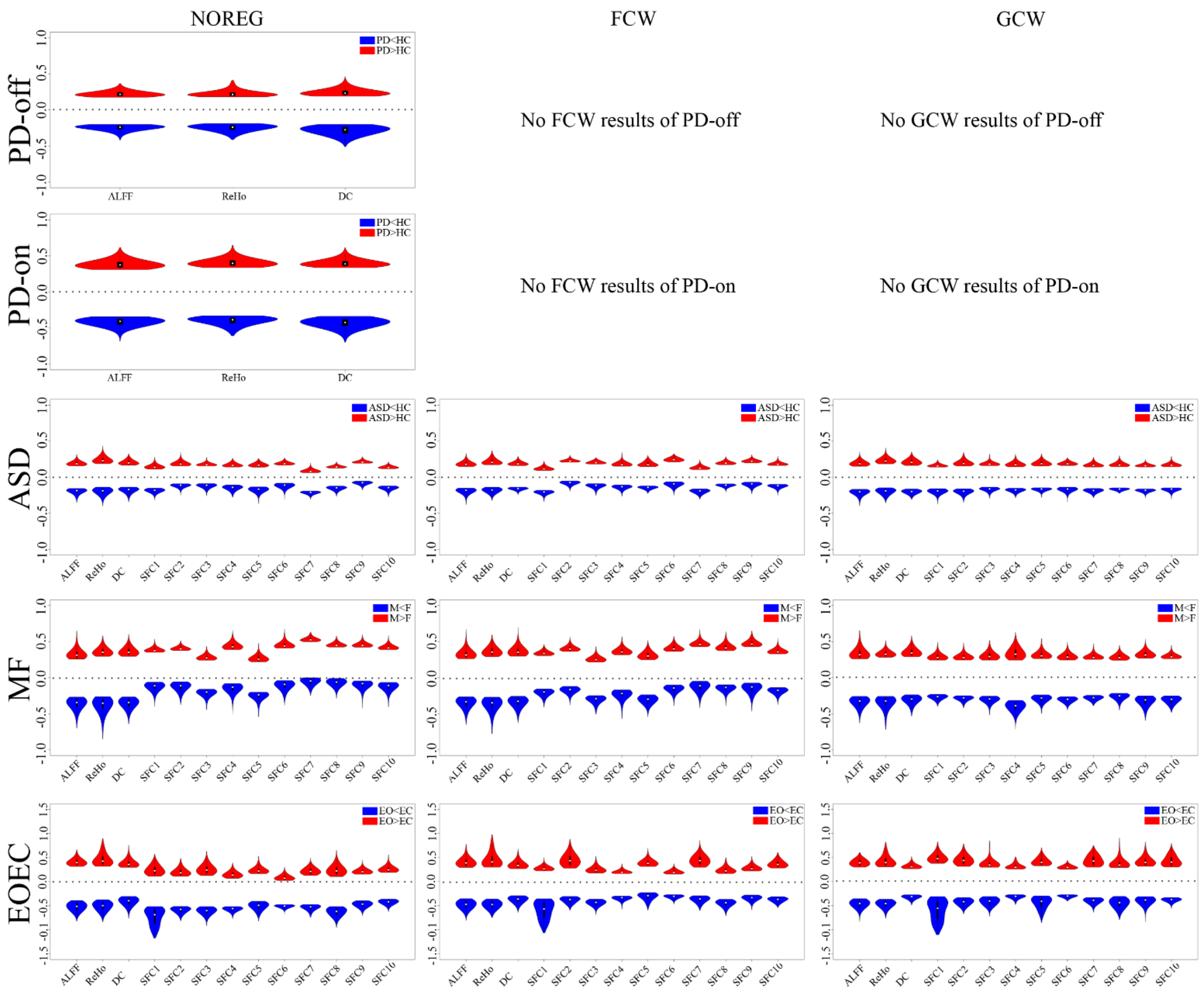

The effect size value (Hedges' *g*) of each voxel was extracted from meta-analytic results. The white dots indicate the median effect size and the black bar indicates the interquartile range of all voxels above threshold for each dataset. X and y axes are metrics and effect size, respectively. PD-off, Parkinson's disease off levodopa vs. healthy controls (HC). PD-on, PD on levodopa vs. HC. ASD, autism spectrum disorder vs HC. MF, healthy male vs. female. EOEC, eye open vs. eyes closed. ALFF, Amplitude of low frequency fluctuation. ReHo, Regional homogeneity. DC, Degree centrality. SFC, Seed-based functional connectivity. NOREG, not regress out covariates in preprocessing. FCW, Regressing out covariates of Friston-24, Cerebrospinal fluid signal, and White matter signal in preprocessing. GCW, regressing out covariates of Global mean time courses, Cerebrospinal fluid signal, and White matter signal in preprocessing. Because of ethical issues, SFC was not analyzed for PD-off and PD-on. It also should be noted that the analysis of regressing out of covariates was not performed in the early stage of data analysis on ALFF, ReHo, and DC because few previous studies did that on the 3 metrics. However, in the late stage, a few co-authors suggested to add regressing out covariates on ALFF, ReHo, and DC as did for SFC. Therefore, regressing out covariates were performed for ASD, FM, and EOEC, but not for PD-off and PD-on. R package vioplot 0.2 was used for plot (https://cran.r-project.org/web/packages/vioplot/index.html).

**Supplementary Figure 6. FNR, accuracy, and FDR for PD-off dataset.**

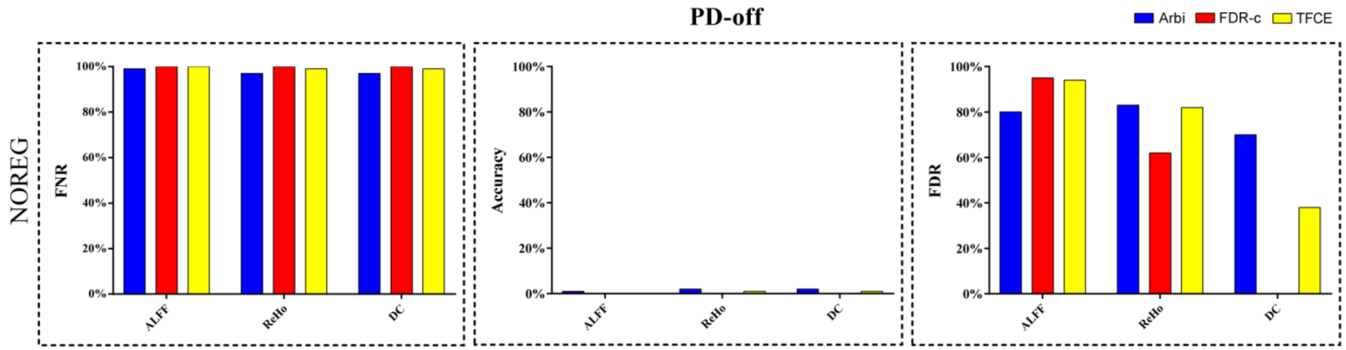


Using meta-analytic results as robust result, the FNR, accuracy, and FDR were calculated for thresholded *t*-image of each study for PD-off datasets. FDR was calculated on those studies which have voxels survived the correction, i.e., if no any voxel survived the correction, FDR was not applicable. Because of ethical issues, SFC was not analyzed for PD-off and PD-on. It also should be noted that the analysis of regressing out of covariates was added in the late stage of data analysis. Either due to the ethical issues, regressing out covariates were not performed for ALFF, ReHo, and DC for PD-off and PD-on. PD-off, Parkinson's disease off levodopa vs. healthy controls. NOREG, not regress out covariates in preprocessing. FNR, false negative rate. FDR, false discovery rate. ALFF, Amplitude of low frequency fluctuation. ReHo, Regional homogeneity. DC, Degree centrality. SFC, Seed-based functional connectivity. GraphPad Prism 7.00 was used for plotting (http://www.graphpad.com).
* FDR was not applicable because no any voxel survived the correction.

**Supplementary Figure 7. FNR, accuracy, and FDR for PD-on dataset.**

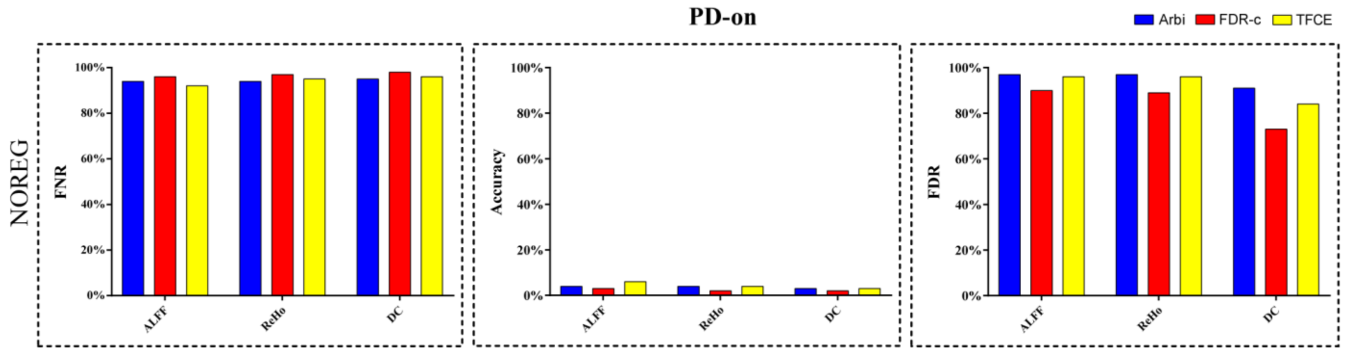


Using meta-analytic results as robust result, the FNR, accuracy, and FDR were calculated for thresholded *t*-image of each study for PD-on datasets. PD-on, Parkinson's disease on levodopa vs. healthy controls. NOREG, not regress out covariates in preprocessing. FNR, false negative rate. FDR, false discovery rate. ALFF, Amplitude of low frequency fluctuation. ReHo, Regional homogeneity. DC, Degree centrality. GraphPad Prism 7.00 was used for plotting (http://www.graphpad.com). Because of ethical issues, SFC was not analyzed for PD-off and PD-on. It also should be noted that the analysis of regressing out of covariates was added in the late stage of data analysis. Either due to the ethical issues, regressing out covariates were not performed for ALFF, ReHo, and DC for PD-off and PD-on.

**Supplementary Figure 8. FNR, accuracy, and FDR for ASD dataset.**

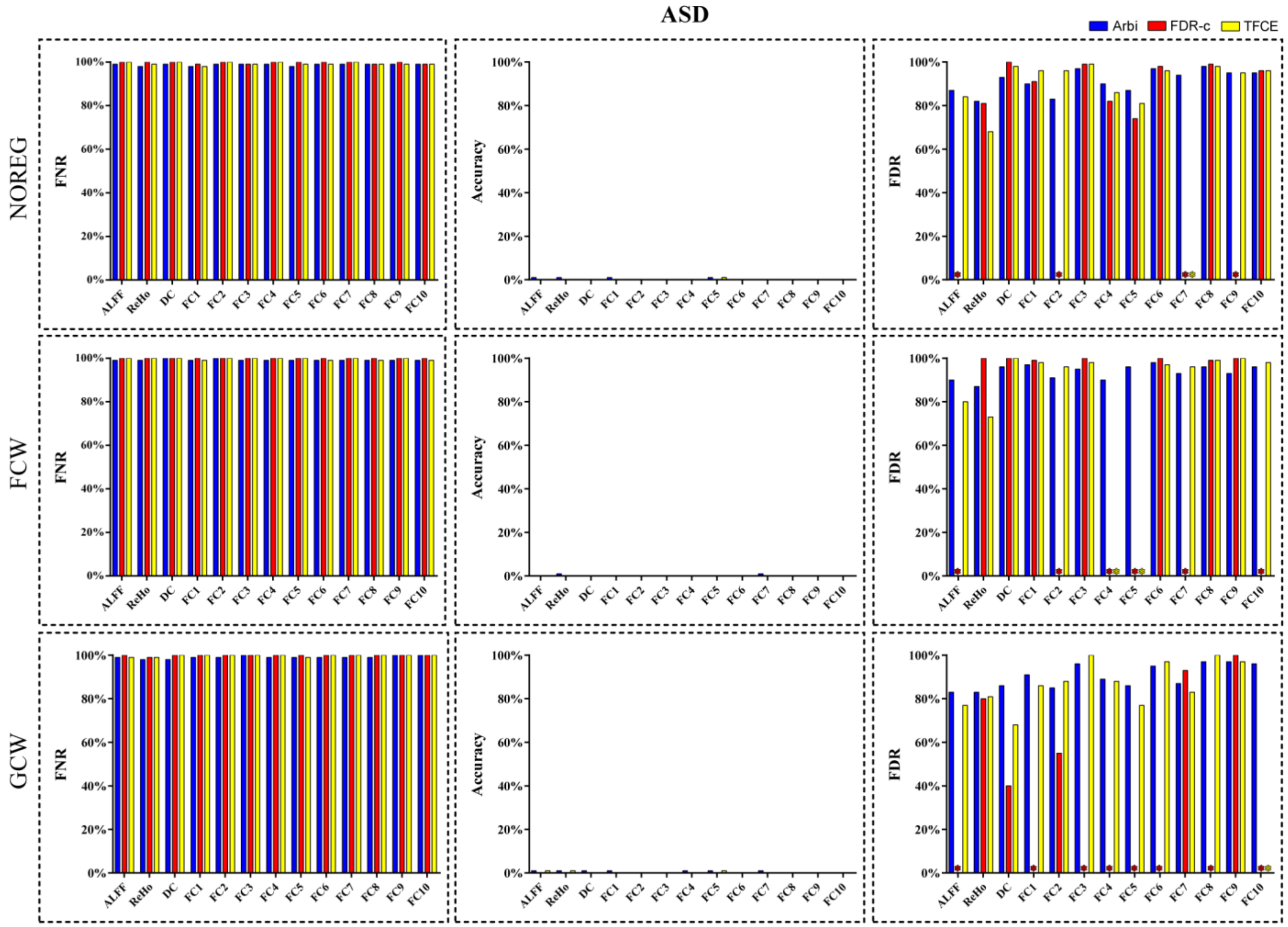


Using meta-analytic results as robust result, the FNR, accuracy, and FDR were calculated for thresholded *t*-image of each study for ASD datasets. FDR was calculated on those studies which have voxels survived the correction, i.e., if no any voxel survived the correction, FDR was not applicable. ASD, autism spectrum disorder vs. healthy controls. NOREG, not regress out covariates in preprocessing. FCW, Regressing out covariates of Friston-24, Cerebrospinal fluid signal, and White matter signal in preprocessing. GCW, regressing out covariates of Global mean time course, Cerebrospinal fluid signal, and White matter signal in preprocessing. FNR, false negative rate. FDR, false discovery rate. ALFF, Amplitude of low frequency fluctuation. ReHo, Regional homogeneity. DC, Degree centrality. SFC, Seed-based functional connectivity. GraphPad Prism 7.00 was used for plotting (http://www.graphpad.com).

* FDR was not applicable because no any voxel survived the correction.

**Supplementary Figure 9. FNR, accuracy, and FDR for MF dataset.**

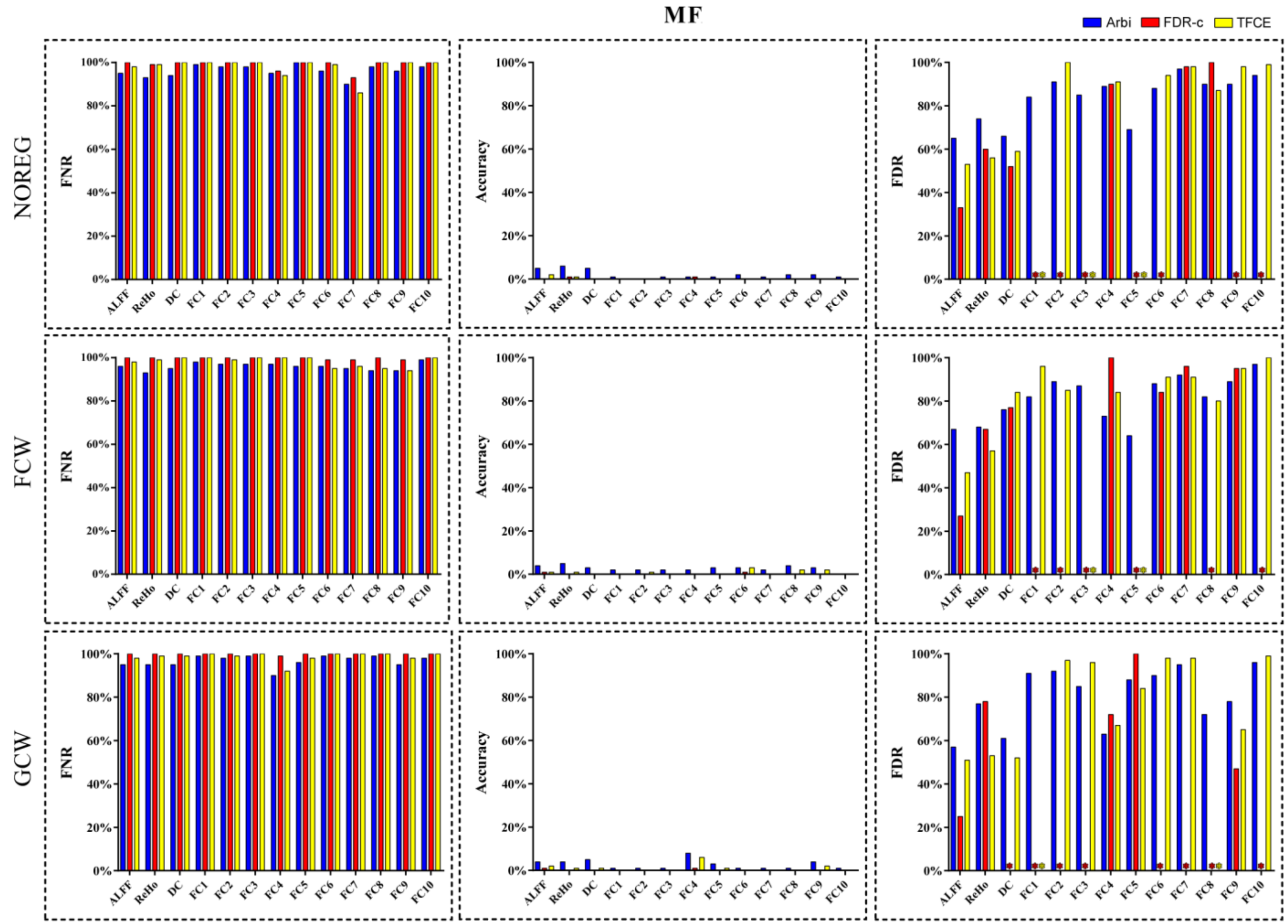


Using meta-analytic results as robust result, the FNR, accuracy, and FDR were calculated for thresholded *t*-image of each study for MF datasets. FDR was calculated on those studies which have voxels survived the correction, i.e., if no any voxel survived the correction, FDR was not applicable. MF, healthy male vs. female. NOREG, not regress out covariates in preprocessing. FCW, Regressing out covariates of Friston-24, Cerebrospinal fluid signal, and White matter signal in preprocessing. GCW, regressing out covariates of Global mean time courses, Cerebrospinal fluid signal, and White matter signal in preprocessing. FNR, false negative rate. FDR, false discovery rate. ALFF, Amplitude of low frequency fluctuation. ReHo, Regional homogeneity. DC, Degree centrality. SFC, Seed-based functional connectivity. GraphPad Prism 7.00 was used for plotting (http://www.graphpad.com).

* FDR was not applicable because no any voxel survived the correction.

**Supplementary Figure 10. FNR, accuracy, and FDR for EOEC dataset.**

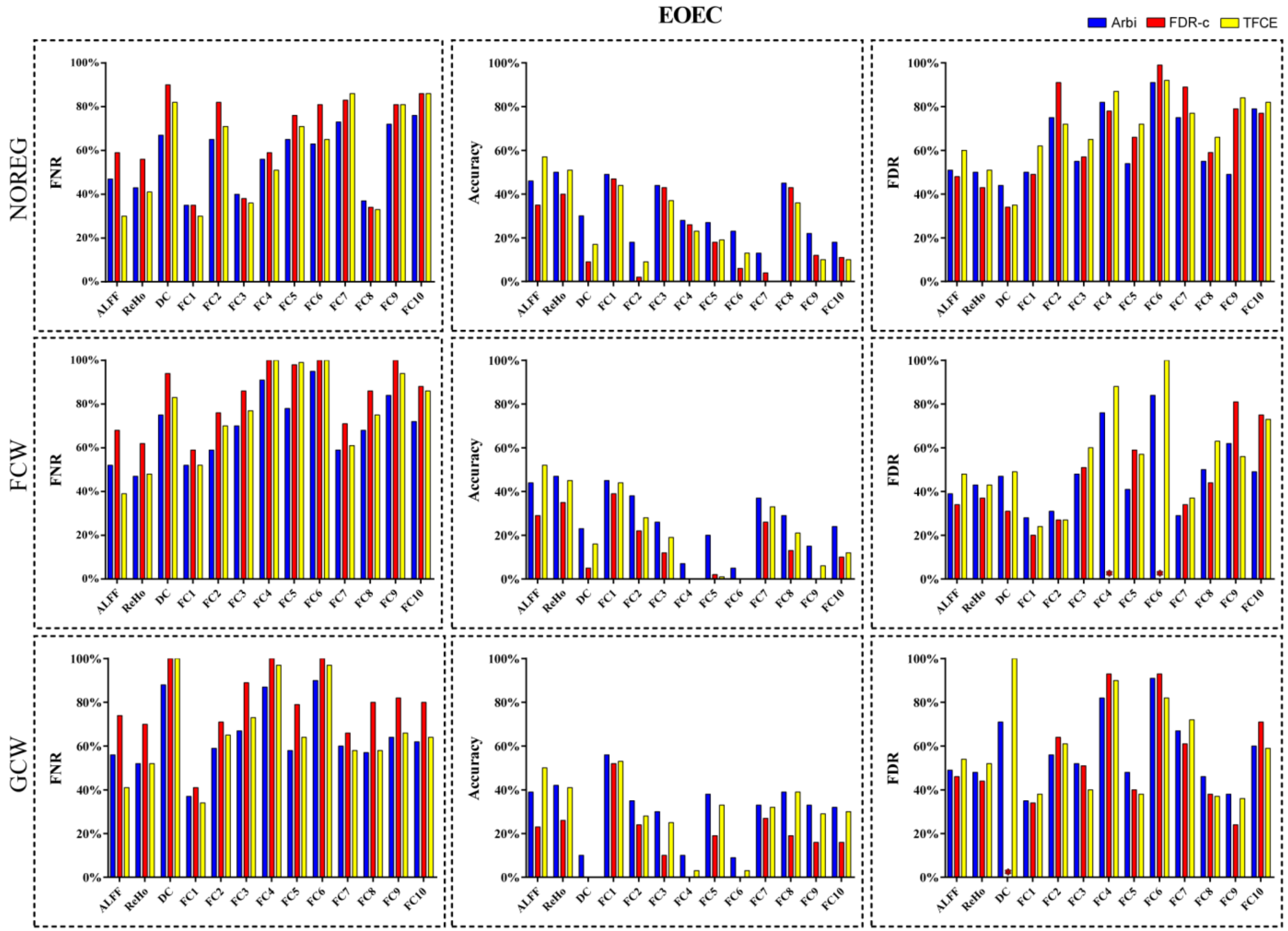


Using meta-analytic results as robust result, the FNR, accuracy, and FDR were calculated for thresholded *t*-image of each study for EOEC datasets. FDR was calculated on those studies which have voxels survived the correction, i.e., if no any voxel survived the correction, FDR was not applicable. EOEC, eye open vs. eyes closed. NOREG, not regress out covariates in preprocessing. FCW, Regressing out covariates of Friston-24, Cerebrospinal fluid signal, and White matter signal in preprocessing. GCW, regressing out covariates of Global mean time courses, Cerebrospinal fluid signal, and White matter signal in preprocessing. FNR, false negative rate. FDR, false discovery rate. ALFF, Amplitude of low frequency fluctuation. ReHo, Regional homogeneity. DC, Degree centrality. SFC, Seed-based functional connectivity. GraphPad Prism 7.00 was used for plotting (http://www.graphpad.com).

* FDR was not applicable because no any voxel survived the correction.
